# SurvBoard: standardized benchmarking for multi-omics cancer survival models

**DOI:** 10.1101/2022.11.18.517043

**Authors:** David Wissel, Nikita Janakarajan, Aayush Grover, Enrico Toniato, María Rodríguez Martínez, Valentina Boeva

## Abstract

Multi-omics data, which include genomic, transcriptomic, epigenetic, and proteomic data, are gaining increasing importance for determining the clinical outcomes of cancer patients. Several recent studies have evaluated various multi-modal integration strategies for cancer survival prediction, highlighting the need for standardizing model performance results. Addressing this issue, we introduce SurvBoard, a benchmark framework that standardizes key experimental design choices. SurvBoard enables comparisons between single-cancer and pan-cancer data models and assesses the benefits of using patient data with missing modalities. We also address common pitfalls in preprocessing and validating multi-omics cancer survival models. We apply SurvBoard to several exemplary use cases, further confirming that statistical models tend to outperform deep learning methods, especially for metrics measuring survival function calibration. Moreover, most models exhibit better performance when trained in a pan-cancer context and can benefit from leveraging samples for which data of some omics modalities are missing. We provide a web service for model evaluation and to make our benchmark results easily accessible and viewable: https://www.survboard.science/. All code is available on GitHub: https://github.com/BoevaLab/survboard/. All benchmark outputs are available on Zenodo: https://zenodo.org/records/11066227.

Key Messages

- We introduce SurvBoard, a comprehensive benchmarking framework for the standardized evaluation of multi-omics cancer survival models. SurvBoard provides an easily accessible platform for the reproducible comparison of models trained on single-cancer and pan-cancer datasets. The platform addresses issues such as the impact of missing modalities and variability in experimental setups. SurvBoard integrates data from four major cancer programs–TCGA, ICGC, TARGET, and METABRIC–to ensure a comprehensive evaluation across diverse types of cancer and research centers.
- SurvBoard results confirm that statistical models generally outperform deep learning models in survival function calibration. We also find that pan-cancer training enhances model performance and that models benefit from incorporating data with missing modalities.
- SurvBoard includes a web service that allows researchers to submit models for benchmarking and evaluation. A leaderboard is accessible via https://survboard.science/ to promote transparency and the continuous assessment of models’ performance.

## Introduction

Survival analysis models for cancer research aim to predict survival-related information using data with censored and truncated observations [Klein et al., 2003]. These models play a crucial role in patient risk stratification and enhancing treatment selection, and are gaining increased interest from both machine learning and bioinformatics communities [Depuydt et al., 2018, Kvamme et al., 2019, Zhong et al., 2021, Tang et al., 2022, Lee et al., 2018].

With the advent of large-scale cancer programs such as The Cancer Genome Atlas (TCGA), International Cancer Genome Consortium (ICGC), and Therapeutically Applicable Research To Generate Effective Treatments (TARGET), researchers have begun to incorporate multi-modal omics data into their survival models [Tomczak et al., 2015, Consortium et al., 2010, Ma et al., 2018]. However, to date, most works have exclusively exploited the TCGA datasets due to their large size and extensive omics information, potentially increasing the risk of overfitting models to this cancer program [Hornung and Wright, 2019, Herrmann et al., 2021, Vale-Silva and Rohr, 2021, Wissel et al., 2023, Cheerla and Gevaert, 2019]. The comparison of survival prediction methods across large-scale cancer programs has also been increasingly difficult due to the many diverging choices regarding data imputation, cancer types under consideration, test splits, and omics types utilized. Furthermore, while a considerable number of benchmarks have explored statistical and regression models for multi-omics integration in the cancer survival context [Herrmann et al., 2021, Zhao et al., 2015, Bøvelstad et al., 2009], no comprehensive benchmark has compared neural and statistical models specifically in the multiomics setting, except our recent work, which was focused almost exclusively on the noise resistance properties of different models, as opposed to overall performance [Wissel et al., 2023].

In a pioneering study, Zhao et al. [2015] benchmarked several feature selection and dimensionality reduction methods combined with the Cox proportional hazards model on four cancer types from the TCGA program. Although there was high variability across cancer types, the study concluded that modalities beyond clinical and gene expression did not significantly enhance prediction performance. However, this study was conducted early in the life cycle of TCGA, and it only considered a limited number of datasets and techniques. More recently, Herrmann et al. [2021] evaluated the performance of 12 statistical multi-omics models in predicting cancer survival across 18 TCGA cancer types. The study found that while incorporating the multi-modal group structure of multi-omics data resulted in better predictions, even the best-performing multi-omics models did not significantly outperform a baseline model trained solely on clinical data. It should be noted that this study excluded neural network models, now frequently employed in survival analysis [Cheerla and Gevaert, 2019, Vale-Silva and Rohr, 2021]. Furthermore, this study did not consider missing modalities or pan-cancer scenarios in the training data, which are increasingly typical in neural networks designed for cancer survival prediction [Cheerla and Gevaert, 2019, Vale-Silva and Rohr, 2021, Fan et al., 2023]. Hornung et al. [2023] provided a comprehensive review and benchmarked multiple methods designed to handle this type of data for a classification task on TCGA. Nießl et al. [2022] used the benchmark design of Herrmann et al. [2021] to illustrate the multiplicity of design options available for benchmarking multi-omics survival analysis. They showed that benchmark results could vary widely depending on the metrics, datasets, and models used.

In addition to different training scenarios and the absence of deep learning models in previous benchmarks, there exist few guidelines for benchmarking survival models more generally. The lack of a standardized benchmarking and experimental framework may cause overly optimistic results due to inadvertent data leakage and the numerous preprocessing options available to researchers when comparing different survival prediction methods [Nießl et al., 2022, Kapoor and Narayanan, 2023].

To address the current gaps in the performance evaluation of multi-omics cancer survival models and to standardize their empirical comparison, we introduce SurvBoard, a comprehensive benchmarking framework. Using SurvBoard, we evaluate the predictive performance of deep learning and state-of-the-art statistical models on datasets from four cancer programs: TCGA, ICGC, TARGET and Molecular Taxonomy of Breast Cancer International ConsortiumMolecular Taxonomy of Breast Cancer International Consortium (METABRIC). SurvBoard allows users to train models in three different settings: standard survival analysis, survival analysis with samples for which some data modalities are missing, and pan-cancer analysis, where a model is jointly trained on multiple cancer types. We showcase the potential use of the SurvBoard platform and discuss common pitfalls in creating datasets for omics survival analysis studies using relevant examples from our four considered cancer programs. Finally, we offer a free web service that displays a leaderboard for SurvBoard, making it easy for other researchers to compare their methods with existing ones.

Going forward, we will restrict ourselves to right-censoring with no truncation, which is typical of most large-scale observational cancer studies and datasets.

## Methods

### Datasets

The SurvBoard benchmark includes a total of 28 cancer datasets from four projects: TCGA, which is arguably the largest and most commonly used database for multi-omics cancer survival analysis (*n* = 21), ICGC, which encompasses and complements TCGA with additional samples from non-American studies (*n* = 4), the pediatric cancer database TARGET (*n* = 2), and the large breast cancer dataset METABRIC (*n* = 1) (Supplementary Table S1, Supplementary Table S2). All datasets from the cancer programs were preprocessed based on the selection criteria highlighted in the Preprocessing section and Supplementary Methods.

### Survival analysis models evaluated in the leaderboard

We evaluated six different approaches on SurvBoard to jumpstart the leaderboard, including two statistical methods and four deep learning models. Our selection of methods was based on the research conducted by Herrmann et al. [2021] and Wissel et al. [2023], who identified BlockForest [Hornung and Wright, 2019] and PriorityLasso [Klau et al., 2018] as the leading statistical methods for accurately predicting clinical outcomes on TCGA datasets.

From previous research, among various multi-modal deep learning architectures for multi-omics survival analysis, the most effective ones were architectures based on the late fusion using an arithmetic mean and intermediate fusion using concatenation [Wissel et al., 2023]. Furthermore, we used two loss functions for the deep learning methods: the commonly used negative logarithm of the Cox PH partial likelihood and the Extended Hazards likelihood, which was recently introduced in a deep learning setting [Zhong et al., 2021, Tseng and Shu, 2011]. We only considered methods that take into account the group structure of the multi-omics data as they have been proven to be more effective than those that do not [Herrmann et al., 2021, Wissel et al., 2023].

Thus, to seed the SurvBoard leaderboard, we conducted experiments for the six methods abbreviated as:

1. **PriorityLasso L1+L2** (with the Elastic-net regularization), a method that orders the input modalities and sequentially uses Elastic-net-based models per modality that are carried forward via offsets into the model fit for the next modality [Klau et al., 2018];
2. **BlockForest**, a method based on random survival forests, that takes the group structure of multi-omics data into account by sampling covariates per modality (as opposed to uniformly) and considers block-specific weights when calculating the split criterion [Hornung and Wright, 2019];
3. Neural network using late fusion with an arithmetic mean and with the Cox PH likelihood, **NN Cox LM** [Ching et al., 2018, Katzman et al., 2018];
4. Neural network using late fusion with arithmetic mean and the Extended Hazards likelihood, **NN EH LM** [Zhong et al., 2021];
5. Neural network using intermediate fusion with concatenation and the Cox PH likelihood, **NN Cox IC** [Ching et al., 2018, Katzman et al., 2018];
6. Neural network using intermediate fusion with concatenation and the Extended Hazards likelihood, **NN EH IC** [Zhong et al., 2021].

Further details regarding the considered models, including hyperparameter choices, can be found in Supplementary Methods.

### Considered modalities

The performance of each model was assessed in three different scenarios: (*i*) on each modality individually, (*ii*) with clinical and gene expression data combined, and (*iii*) with all modalities available for that particular dataset. Notably, available modalities significantly varied across cancer programs and datasets (Supplementary Table S1).

Furthermore, for experiments where only one modality was used and no multi-modal integration was required, equivalent models that did not take group structure into account were employed. For example, Elastic Net was used instead of PriorityLasso, and Survival Random Forest instead of BlockForest in the unimodal experiments. For all deep learning models, the unimodal experiments used a standard Multilayer perceptron (MLP).

### Three settings for the evaluation of survival models

SurvBoard allows users to train models in three settings: standard, missing data modality, and pan-cancer.

#### Standard setting

Our first setting implements standard multi-omics survival analysis. Each model is trained and evaluated only on samples of the same cancer type.

#### Missing data modality setting

The missing data modality setting refers to the scenario in which several samples in a dataset lack data for one or more modalities but still have data for some modalities, in addition to survival information. This is common in TCGA, where many patients lack protein expression data. Thus, models that can handle samples with missing modalities benefit from an increased training set size.

In the missing data modality setting of SurvBoard, the tumor samples with data lacking one or more modalities were used as additional training data for models that can handle missing modalities. In the test sets, only those samples with complete data modalities in all settings were present. This was done to ensure comparability with other models by using a consistent test set.

#### Pan-cancer setting

In the pan-cancer setting, we jointly trained models on datasets from different cancer types. However, since not all datasets include all modalities, models that cannot handle missing modalities cannot be trained in the pan-cancer setting when all modalities are used.

Therefore, in our pan-cancer experiments, we only used clinical data and gene expression. This allowed us to obtain pan-cancer results for all models included in SurvBoard.

It is worth noting that the pan-cancer scenario only applies to the TCGA project, as other projects did not provide data that was normalized in a unified way for a pan-cancer analysis.

### Preprocessing

While several packages (*e*.*g*., Cerami et al. [2012]) and data sources (*e*.*g*., Weinstein et al. [2013]) allow the acquisition and usage of TCGA, ICGC, TARGET, and METABRIC datasets, preprocessing choices are left to the user, which leads to inconsistency across benchmarking experiments. To enable a fair comparison of existing and new methods, SurvBoard standardizes most preprocessing choices.

#### Endpoint choice

While all of our considered cancer datasets provide multiple endpoints, *e*.*g*., overall survival (OS), Disease Free Survival (DFS), and others, it is relatively common in the survival literature to utilize OS, as it is ubiquitously available compared to other endpoints. For example, for TCGA, the Clinical Data Resource (CDR) analyzed the suitability of different endpoints for survival analysis and found that OS is the most used and generally appropriate for most datasets [Liu et al., 2018]. The only exception was the situation where progression was the event of interest, in which case the progression-free interval was recommended. We follow this broader convention and use OS as the endpoint of all datasets within our benchmark.

#### Patient cohort

We restricted our datasets to primary tumor tissue samples. We also excluded patients for whom either the event indicator or the event time was missing.

#### Dataset selection

We followed the methodology of Herrmann et al. [2021] and selected only datasets with at least 100 samples and a minimum event ratio of 5% or 10 total events, whichever was larger. This ensured that we could compute meaningful performance metrics. We only counted samples with complete modalities for dataset selection since only these were included in the test splits. Samples with missing modalities were only used as additional training data.

#### Modality selection

We chose the maximum number of modalities available for each dataset in each cancer program and excluded datasets lacking clinical data and gene expression modalities. In addition, we selected only datasets that fulfilled the criteria above for at least two omics modalities, leading to a total minimum of three modalities for each dataset.

#### Clinical variables

To ensure a fair comparison between different cancer programs and cancer types, we only considered standard clinical information that was available at the time of diagnosis to prevent data leakage. This included demographic data such as age and gender and staging variables such as clinical stage. For each cancer program and dataset, we used slightly different variables in the SurvBoard framework (as outlined in Supplementary Methods and Supplementary Table S1). We chose not to include information specific to certain cancer types in our analysis, such as smoking history for lung cancer.

#### Missing values within modalities

To handle missing data, we followed a three-step procedure. Firstly, we created a token for non-available (NA) information for categorical variables. We assumed that the missingness of categorical variables might correlate with either the target variable or other covariates, which is known as Missing Not At Random (MNAR) [Van Buuren, 2018]. Our goal was to avoid mixing unrelated categories, so we did not use mode imputation. Secondly, non-categorical variables missing in more than 10% of samples in a specific dataset were excluded from that dataset. Thirdly, non-categorical variables with missing rates less than 10% in a dataset were imputed using the median of the available samples on the full dataset. Although imputation in the full dataset could lead to some information leakage, previous research has shown that it does not cause significant bias [Hornung et al., 2015]. This approach was designed to eliminate non-model-specific preprocessing choices from researchers.

#### Missing modalities

To create splits for training and testing for each dataset of each cancer program, we created two sets of samples: a “complete” set used in the standard setting and an “incomplete” set that included samples with one or more missing modalities. Importantly, the incomplete set was intended only as additional training data in our benchmark, as noted in the section describing the three training settings. Within the incomplete set, NA values indicated that a particular modality was missing in a specific sample. As explained above, NA values for particular variables within available modalities were no longer present in the incomplete set as they had been imputed or removed.

#### Pan-cancer training

For the TCGA program, which provided data normalized in a pan-cancer way, we combined variables across all cancer types in the pan-cancer dataset. The variables that were not available for all cancer types were excluded. However, if a particular cancer type lacked a modality, we did not remove this modality from the pan-cancer dataset. Instead, we marked it as missing for the samples corresponding to that particular cancer type.

### Performance metrics

We measured three performance metrics in our benchmarks, all of which are evaluated on the survival function level. Firstly, we used Antolini’s Concordance (Antolini’s C) to assess the ability of each survival model to discriminate low-risk patients from high-risk patients over time [Antolini et al., 2005]. Secondly, we evaluated the Integrated Brier score (IBS), which is a widely used measure in survival benchmarks that assesses both discrimination and calibration accuracy [Graf et al., 1999]. Thirdly, we included the recently proposed D-Calibration (D-CAL), which measures the distributional calibration of each multi-omics survival model [Haider et al., 2020]. For D-CAL, we evaluated the test statistic where lower values corresponded to a better fit (Supplementary Methods).

### Validation

We implemented five-fold cross-validation, repeated five times, resulting in a total of 25 test splits for each cancer type. To create the splits, we stratified the data by the event indicator, OS, which ensured that the event ratio in each train and test fold was the same as in the original unsplit data. The incomplete modality samples were not part of the test set and were instead used only as additional training data. In the pan-cancer setting, training data from all cancer types were included in each training split.

In cases where a model encountered numerical issues or sparse methods reported a fully sparse fit as the best model, we used a Kaplan-Meier estimator as a replacement [Kaplan and Meier, 1958]. For instance, the Lasso model has been observed to sometimes fail in very high-dimensional datasets with high multicollinearity due to numerical issues [Herrmann et al., 2021, Sohn et al., 2009]. We note that other choices have been explored here and that this choice may have an impact on results [Herrmann et al., 2021, Nießl et al., 2022]. Despite this, to enable a fair comparison and prevent gameability for future submissions to SurvBoard (for example, by deliberately setting difficult splits to failures), we settled on the choice of a simple Kaplan-Meier replacement (Supplementary Methods, Supplementary Table S3).

## Results

### Benchmark design

We developed a benchmark framework named SurvBoard that allows for the thorough evaluation of multi-omics survival models in the context of cancer. Our framework, SurvBoard, has several unique features (Fig. 1). Firstly, we used datasets from four different cancer programs, TCGA, ICGC, TARGET, and METABRIC, containing data from up to seven modalities including clinical variables, gene expression, somatic mutations, DNA methylation, copy number alterations, protein expression from reverse-phase protein arrays (RPPA), and miRNA expression (Supplementary Table S1 and Supplementary Table S2, Methods, Supplementary Methods).

**Fig. 1.**
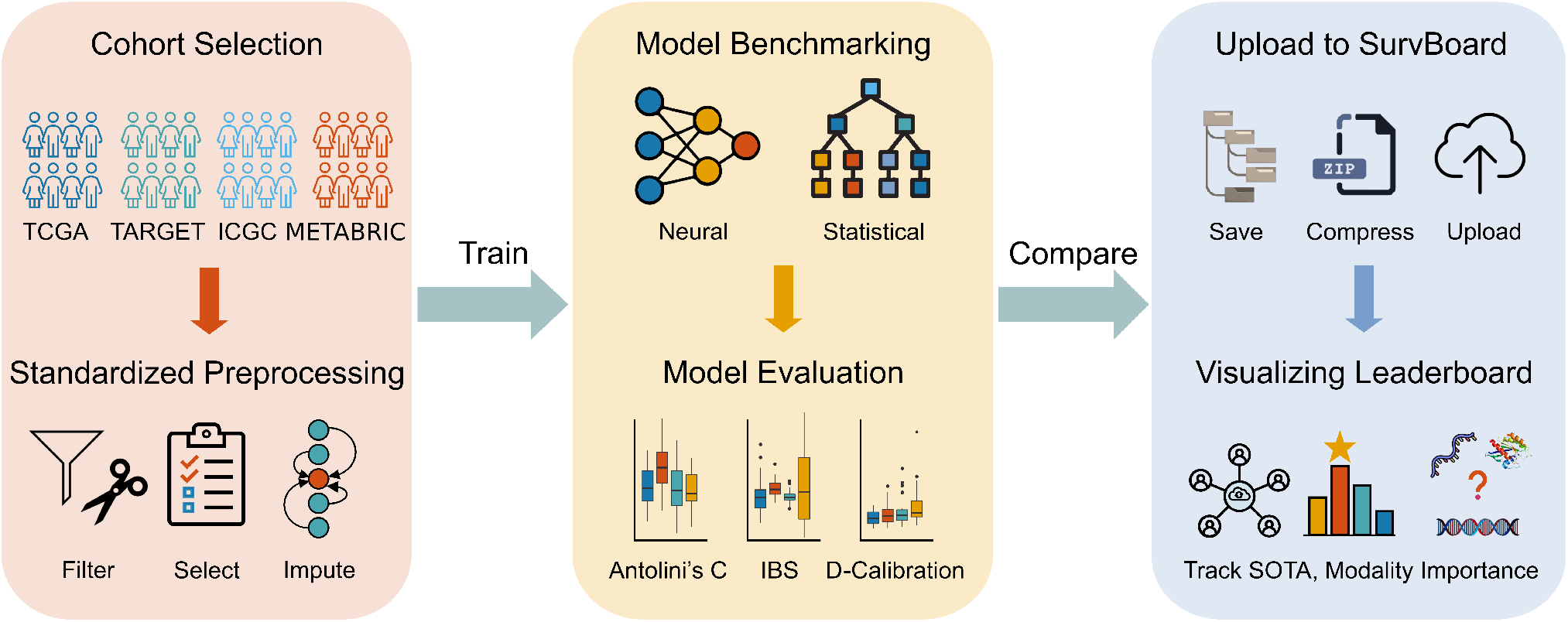
The SurvBoard framework enables the reproducible, easily accessible, and standardized comparison of (multi-)omics cancer survival methods. SurvBoard is based on a careful cohort selection from four cancer programs: TCGA, ICGC, TARGET, and METABRIC. The datasets from all programs are preprocessed in a standardized manner, which allows for uniform assessment of the models created. The evaluation results can then be uploaded to the SurvBoard leaderboard to track model results.

Secondly, the datasets were filtered and preprocessed in a standardized manner to enable optimal comparability across models. Stratified splits were created to facilitate the uniform assessment of model performances (Methods).

Thirdly, we prepared the datasets for conducting experiments in three different settings: (*i*) a standard setting, where all samples present all modalities used in training and the model training is performed individually for each cancer type, (*ii*) a setting where certain patients do not have information for specific modalities, and (*iii*) a pan-cancer training setting where multiple cancer types are trained jointly via a unified model (Methods).

Last, we assessed the model performance using three different metrics that focus on the accuracy of patient outcome prediction and model calibration: Antolini’s C, IBS, and D-CAL (Methods, Supplementary Methods, Supplementary Table S4).

### Leaderboard

Additionally, we have developed a web service that enables researchers and other stakeholders to submit predictions on the SurvBoard benchmark set, which can be accessed via https://www.survboard.science/. Using this service, one can also download and inspect previous submissions, including the provided baselines. SurvBoard’s web service evaluates submitted predictions and displays the performance metrics for all datasets within the benchmark in an easy-to-compare leaderboard format (Fig. 2 and Supplementary Fig. S1-S6). To access the sample submission file and links to our web service, please visit the GitHub repository at https://github.com/BoevaLab/survboard/.

**Fig. 2.**
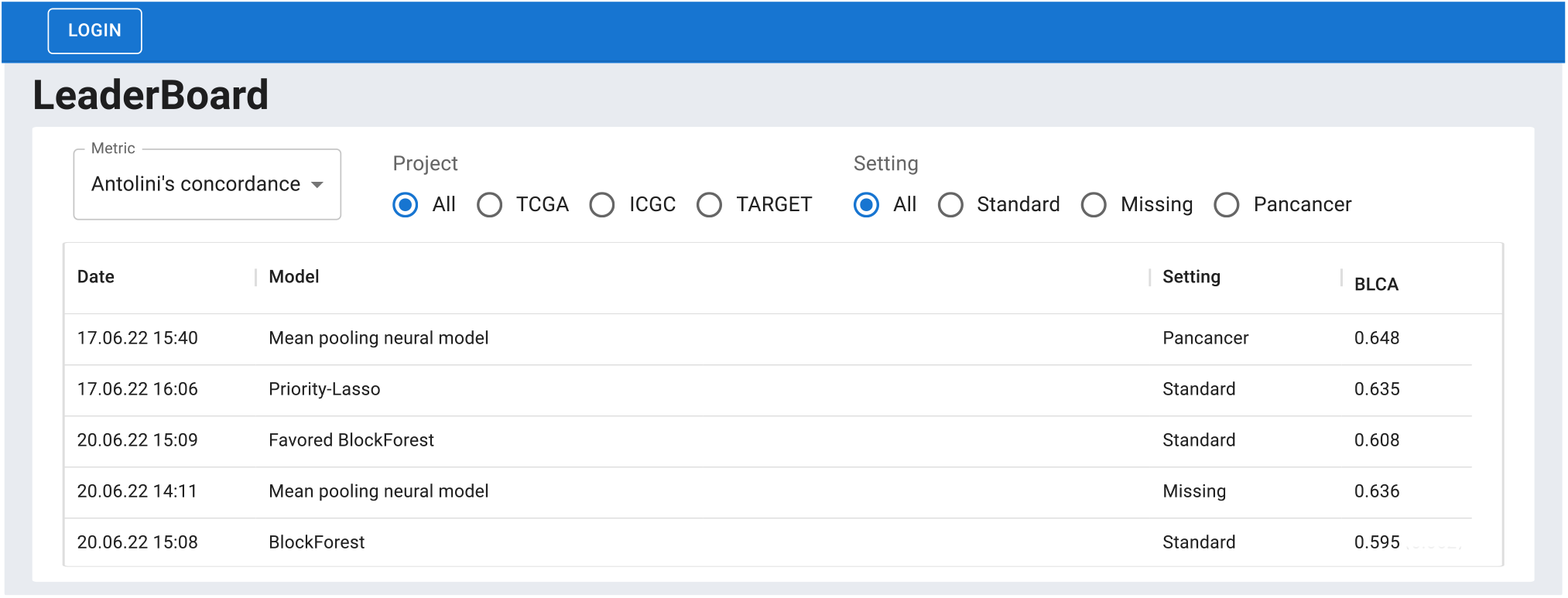
The SurvBoard web service curates and makes model results submitted to SurvBoard easily explorable and downloadable. The web service also ensures that SurvBoard stays up to date, as other researchers can easily extend our initial baseline models.

We seeded SurvBoard by submitting six models: two statistical and four deep learning models trained on various combinations of input modalities and in different settings (Methods). We limited the selection of models to those that had already demonstrated top performance in multi-omics cancer datasets in previous benchmarks [Herrmann et al., 2021, Wissel et al., 2023] (Methods).

### Assessment of model performances

To be fair to each model and to evaluate overall survival prediction performance, we first determined on which combination of modalities each model performed the best. We trained each model on each available modality unimodally, clinical data and gene expression together (which has performed well in related work), and all available modalities together. We selected the best modality set based on Antolini’s C metric for each model (Methods). We found that clinical variables and gene expression data were the most predictive modalities across all models (Supplementary Fig. S7).

Next, we evaluated the performance of each model on their optimized modality sets using three performance metrics: Antolini’s C, IBS, and D-CAL. We observed that overall, BlockForest trained on clinical variables and gene expression data performed the best among all models (Fig. 3A-C). Notably, BlockForest achieved the best rank across datasets for the IBS and performed second best in terms of both D-CAL and Antolini’s C. Prioritylasso L1+L2 also performed well, achieving the best rank for D-CAL and the second-best rank for the IBS, while performance for Antolini’s C was generally more variable and no clear winner emerged.

**Fig. 3.**
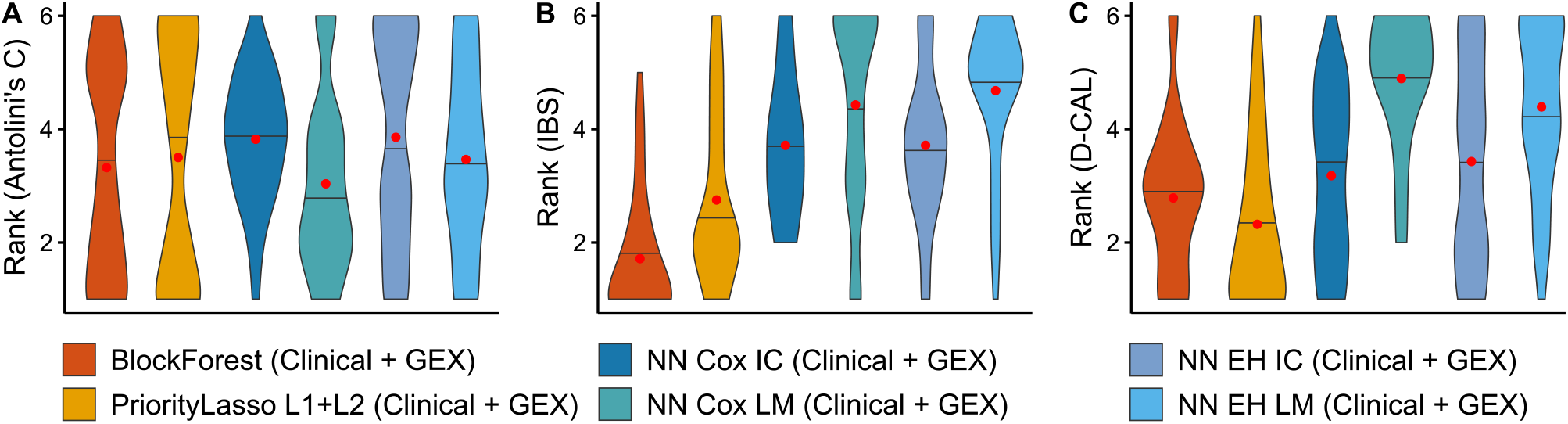
Regularized linear models and random-forest-based methods outperformed deep learning methods on the SurvBoard benchmark. Overall ranked results across all datasets, where lower ranks indicate better scores. **A**. Antolini’s concordance (Antolini’s C). **B**. Integrated Brier Score (IBS). **C**. D-Calibration (D-CAL). Each model was trained on the dataset on which it ranked the best among itself in terms of the median Antolini’s C rank (see Figure S2). Ties in the median Antolini’s C rank were broken using the mean Antolini’s C rank. The median is indicated by black horizontal lines, the arithmetic mean by red points.

Deep learning methods generally could not compete with the statistical models PriorityLasso L1+L2 and BlockForest. While NN Cox LM performed well for Antolini’s C, achieving the best rank, it performed poorly in terms of model calibration, scoring the worst median rank for both the IBS and D-CAL. Other deep learning-based methods performed similarly, achieving overall worse results than PriorityLasso L1+L2 and BlockForest in terms of IBS and D-CAL, while achieving variable results on Antolini’s C, with three out of four deep learning methods outperforming PriorityLasso L1+L2.

It is worth noting that the best model configurations in terms of the IBS metric differed from those selected based on Antolini’s C (Supplementary Fig. S8). Most models achieved optimal IBS values when only clinical variables were used for predictions. However, even with using the input modalities that led to the best IBS for each model (Supplementary Fig. S8), the results were generally consistent with those obtained by optimizing Antolini’s C metric (Supplementary Fig. S9). Nevertheless, when using only clinical data for training, the survival problem becomes significantly more straightforward and the integration becomes unnecessary, hence leading to little or no differences between some of the deep learning methods.

In addition, discriminative survival model prediction performance as measured by Antolini’s C was noticeably more concordant between deep learning methods than between PriorityLasso L1+L2 and BlockForest or either of the former and any of the deep learning methods (Supplementary Fig. S10).

### Added value of the pan-cancer training and including additional training samples with missing modalities

Using the SurvBoard framework, we determined to what extent pan-cancer training, *i*.*e*., (*i*.*e*.,) simultaneous training on all datasets from a cancer program, could improve the performance of omics survival analysis models. We used the two most informative modalities, clinical variables, and gene expression data, as input for the assessment. The results showed that pan-cancer training improved the median performance for most considered methods in terms of Antolini’s C and the IBS, while the impact on D-CAL was much more variable. Moreover, the performance increase was often statistically significant (Fig. 4A). Deep learning methods benefited the most from pan-cancer training, with all considered neural network models significantly improving their performance for at least two out of the three considered metrics. Interestingly, however, the best-performing method in the standard setting, BlockForest, benefited the least from pan-cancer training, with the D-CAL metric getting significantly worse.

**Fig. 4.**
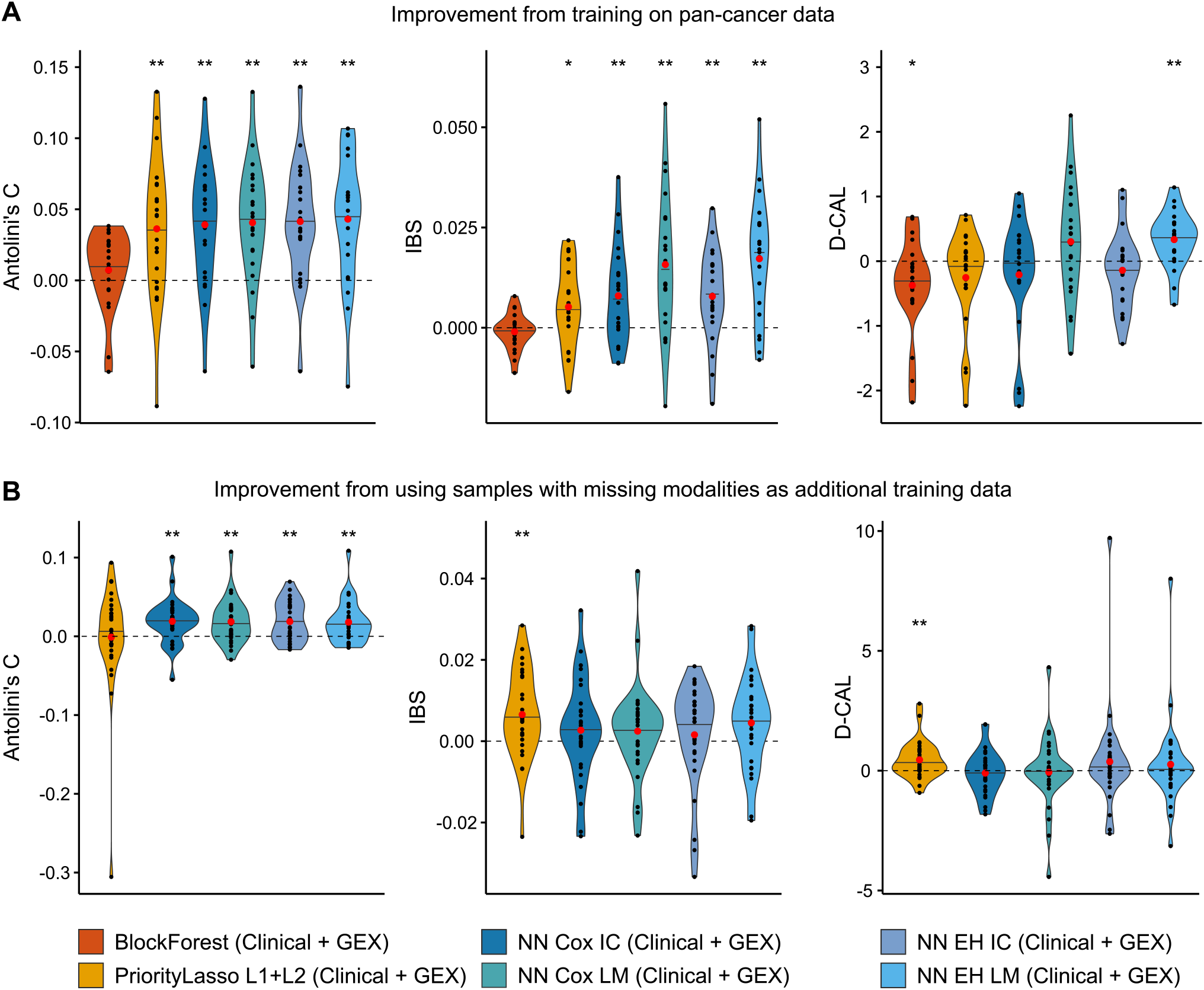
Training on multiple cancer types jointly (pan-cancer) and leveraging samples with some missing modalities improved performance for most models and most metrics. **A**. Pan-cancer training based on clinical and gene expression data improved all three considered metrics for most models, often significantly. **B**. Training on all available modalities that also included samples with missing modalities on improved Antolini’s C for all deep learning methods and both the IBS and D-CAL for PriorityLasso, relative to training on all available modalities without missing modality samples. Significance indicated by * (*p* < 0.05) and ** (*p* < 0.01), based on a two-sided Wilcoxon signed-rank test. The median is indicated by black horizontal lines, and the arithmetic mean by red points.

Next, we investigated to what extent including samples with some missing modalities during training could improve model prediction performance on unseen samples with all modalities present. For this, we performed experiments on all models capable of handling missing modalities (Supplementary Methods), namely all models except BlockForest. Since the clinical data and gene expression setting had no missing modality samples, we considered all available modalities and compared the performance of each method with and without including missing modality samples as additional training data. After including into the training set samples with missing modalities, Antolini’s concordance improved significantly relative to the non-missing modality models for all considered deep learning methods but not for PriorityLasso L1+L2 (Fig. 4B). Meanwhile, only PriorityLasso L1+L2 showed significant improvement in model calibration as measured by IBS and D-CAL (Fig. 4).

### Take-aways for effective model development and validation

In our benchmark framework, we have aimed to remedy potential pitfalls related to the training and validation of omics cancer survival models. The pitfalls discussed below emphasize the importance of some of the design choices we made in the SurvBoard benchmark and may be helpful for other researchers to validate their models on small *n* and large *p* data size regimes.

First, it is essential to report both discriminative and calibration metrics while evaluating survival models. Discriminative metrics such as Harrell’s concordance and Antolini’s C have been widely used, along with calibration metrics such as the IBS. These metrics do not necessarily correlate, with correlations close to zero or even negative on some datasets (Fig. 5A-B). Thus, reporting at least one metric of each type is crucial. We found that on various datasets included in SurvBoard, models could be favored if only one metric was reported. For example, on the METABRIC breast cancer dataset, PriorityLasso outperformed all other models in terms of the IBS and D-CAL while achieving among the worst concordance values as measured by Antolini’s C out of all methods (Fig. 5B).

**Fig. 5.**
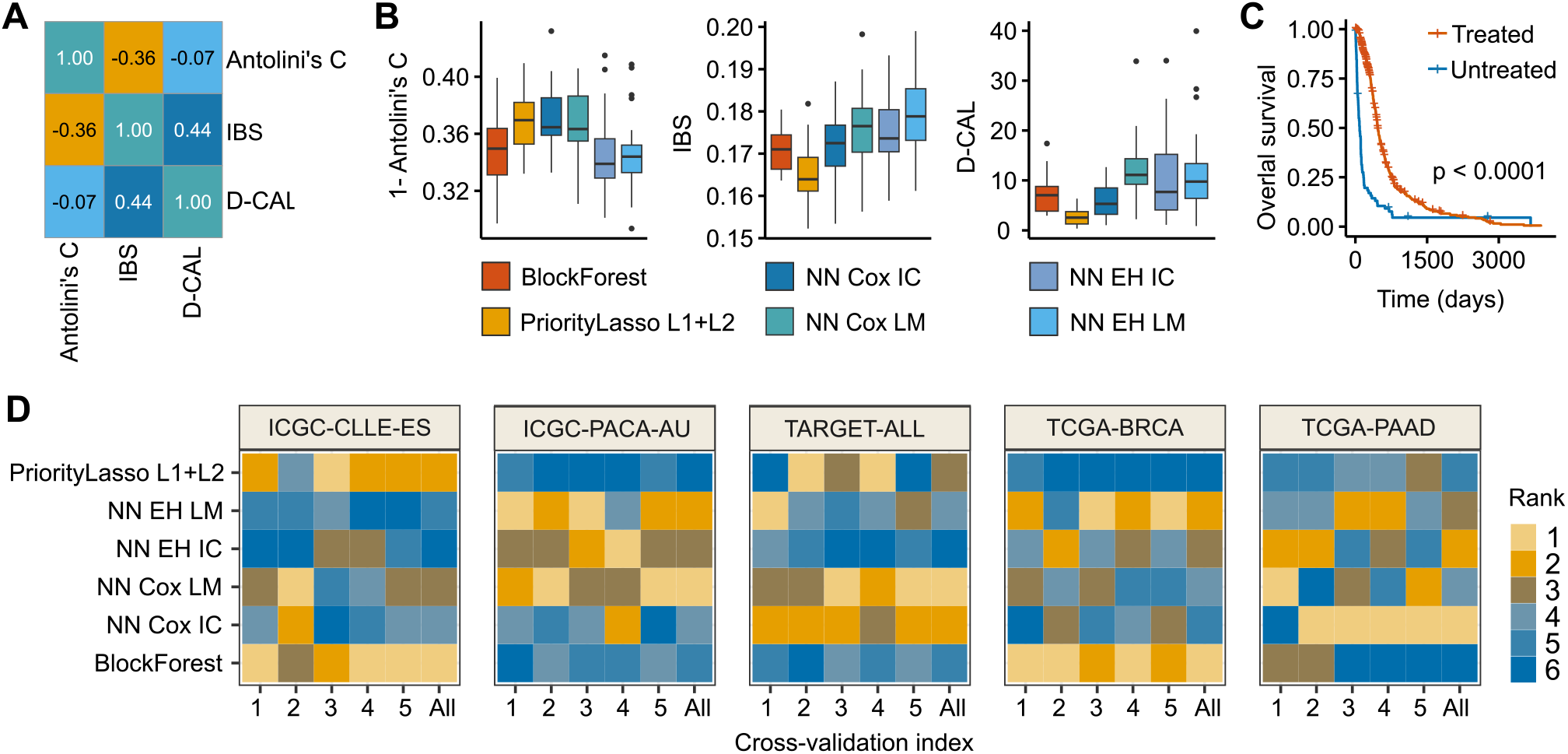
There are several pitfalls to consider when benchmarking and validating (multi-)omics survival models. **A**. The usage of several survival metrics is necessary, since the discriminative and calibration performance of a particular model may not be correlated. Pearson correlation matrix between Antolini’s C, IBS, and D-CAL of all models on breast cancer (METABRIC). **B**. On the METABRIC breast cancer dataset using clinical and gene expression data, PriorityLasso L1+L2 achieves the best performance for both D-CAL and IBS but is among the worst performers for Antolini’s C. Performance of selected multi-omics survival methods on breast cancer (METABRIC) as measured by 1 - Antolini’s concordance, the Integrated Brier Score, and D-Calibration. Lower values are better. **C**. Treatment-related variables should be treated in survival analysis models to avoid any potential data leakage [Kapoor and Narayanan, 2023]. Kaplan-Meier estimator on glioblastoma multiforme (TCGA-GBM) stratified by whether a patient received radiation therapy. Patients with an unknown value for radiation therapy were excluded. **D**. Especially for small datasets, it is essential to use repeated cross-validation to prevent dependence on a particular cross-validation split. Antolini concordance rank of all considered models across cross-validation repetitions on all cancer types; one signifies best rank, five worst rank.

Second, when using multi-omics survival methods, the choice of clinical variables is crucial for ensuring high performance. However, although it is tempting to use all treatment-related and outcome-related covariates as predictive features, including some of these may lead to data leakage [Kapoor and Narayanan, 2023]. For instance, a clinician might decide against starting radiation therapy if the patient is expected to have a short life expectancy due to their illness or other factors [Arenas et al., 2014]. Indeed, on TCGA, we observed cancer types in which treatment-related variables such as “radiation therapy” were strongly associated with the outcome (Fig. 5C), which could be either due to a treatment effect or (partially) an effect of not prescribing the treatment due to a very advanced disease stage. It is thus advisable to be mindful of the choice of clinical variables, especially treatment variables when benchmarking survival prediction methods.

Third, it is advisable to include a large spectrum of studies and datasets as possible in a benchmark to account for variability in model performance. In SurvBoard, we reported results on datasets from four cancer programs. This may guard against overfitting to a particular cancer program that is frequently used in the literature or related work.

Fourth, one should avoid using unrepeated cross-validation or even a single split when reporting model prediction performance since this can make model ranking inconsistent. Indeed, in SurvBoard, we observed large variability in model rankings by Antolini’s C metric across cross-validation repetitions for selected models (Fig. 5D). Smaller datasets, as measured by the number of events *e*, rather than the number of samples *n*, such as the Acute Lymphoblastic Leukemia (TARGET-ALL) dataset, were especially prone to this issue. Performance results on larger datasets tend to show greater consistency but may still suffer from sizeable variability. Thus, we suggest performing several repetitions of cross-validation on each dataset.

Fifth, it is crucial to ensure the comparability of past and future work. For example, when utilizing samples with missing modalities or training models on multiple cancer types, it is imperative to choose train and test splits that can also be utilized by models not utilizing these settings (for example, samples with missing modalities should not be part of the test set since this makes comparison with non-missing modality models impossible). To circumvent this issue, SurvBoard employed missing modality samples as additional training data instead of incorporating them into the test sets.

Finally, in addition to making the code for models publicly available, it is vital to focus on providing a reproducible hyperparameter tuning strategy and evading manual hyperparameter optimization to enhance the model’s reusability by other researchers.

## Discussion

In this work, we presented SurvBoard, a rigorous benchmark and a framework for the validation and comparison of omics survival models. In a proof of concept, here, SurvBoard enabled the comparisons of six models across 28 datasets from four projects. SurvBoard focused on model comparability by ensuring that models utilizing pan-cancer data or samples with missing modalities can be compared to models trained on single datasets. Additionally, we provided a simple web service that allows researchers to evaluate their models on our new benchmark easily. In our work, we also illustrated potential pitfalls during the validation of omics survival models, highlighting the importance of the choice of clinical variables, the use of repeated cross-validation, and the display of results using several relevant performance metrics.

Our observations that statistical models often outperform deep learning ones for the survival prediction in cancer and that clinical variables and gene expression data constitute the two most informative modalities were consistent with our earlier work [Wissel et al., 2023] and the work of Herrmann et al. [2021]; however, the current analysis encompassed a broader array of datasets and cancer programs. We also showed how the SurvBoard benchmarking platform enables novel findings. We demonstrated the positive effect of pan-cancer training for most of the survival analysis models considered in our leaderboard and examined the effect of conducting training on samples with missing data modalities [Cheerla and Gevaert, 2019, Vale-Silva and Rohr, 2021, Fan et al., 2023].

In the future, further investigation is required to explore new model architectures and loss functions, particularly those that have been previously used in clinical survival analysis datasets [Kvamme et al., 2019, Tang et al., 2022]. Methodological research could also focus on the problem of low model resistance to the inclusion of less informative data modalities. Indeed, many methods continued to perform worse in our benchmark when more modalities were added, as evidenced by the fact that few models reached their best performance for any of the metrics when all modalities were included (Supplementary Fig. S7 and Supplementary Fig. S8).

It is important to consider some limitations of our work. While SurvBoard uses datasets from multiple cancer programs, the TCGA program contributed significantly more datasets than the others. Therefore, the conclusions drawn from smaller projects may be less reliable in comparison to those drawn from the larger TCGA dataset. Additionally, certain design decisions made during SurvBoard development may have an impact on the work outcomes. Therefore, results might vary if different design decisions had been made [Nießl et al., 2022]. Nevertheless, we argue that defining a common benchmarking set for multi-omics survival analysis models is crucial to enable comparability across models, even if the benchmark has certain biases. Indeed, several standard computer vision datasets have recently been found to contain label errors and other inaccuracies. Despite these errors, the datasets have greatly contributed to the progress of method development in this field [Northcutt et al., 2021].

Within SurvBoard, we excluded comparisons of runtime and memory requirements since we cannot correctly assess these computational requirements when researchers submit future model predictions to our web service later on in the lifecycle of SurvBoard. Furthermore, to ensure model comparisons in the pan-cancer setting, our datasets only included standard clinical variables such as demographics or staging, which may be a disadvantage to some methods.

To sum up, the development of consistent preprocessing pipelines and online resources for evaluating multi-omics survival models is crucial to advancing research in the field of cancer. In the future, we expect our benchmarking framework to lead to more reliable conclusions about the superiority of different models in predicting patient survival.

## Acknowledgments

The results shown here are in whole or part based upon data generated by the TCGA Research Network: https://www.cancer.gov/tcga. The results shown here are in whole or part based upon data generated by the ICGC: https://dcc.icgc.org. The results published here are in whole or part based upon data generated by the Therapeutically Applicable Research to Generate Effective Treatments (https://ocg.cancer.gov/programs/target) initiative, phs000218.

**Davis Wissel** is a PhD candidate at ETH Zurich, Computer Science Department, Institute for Machine Learning, Zurich, Switzerland, and the Department of Molecular Life Sciences at the University of Zurich, Switzerland. His research interests center around survival analysis methods for omics data as well as methods for and the analysis of long-read RNA-seq data.

**Nikita Janakarajan** is a PhD candidate at IBM Research Europe, Zurich, Switzerland and ETH Zurich, Computer Science Department, Institute for Machine Learning, Zurich, Switzerland. Her research interests lie in multi-modal representation learning, particularly in the context of cancer biology, to gain valuable insights for the development of personalized therapies.

**Aayush Grover** is a PhD candidate at ETH Zurich, Computer Science Department, Institute for Machine Learning, Zurich, Switzerland. His research interests are deep learning and cancer epigenomics.

**Enrico Toniato** is a Technical Leader of the Data and AI Team at IBM Research Europe, Zurich, Switzerland. He specializes in the research and development of large language models for code generation.

**María Rodríguez Martínez** is a professor at Yale School of Medicine, Department of Biomedical Informatics & Data Science, New Haven, Connecticut, USA. Her research focuses on computational immunology, and developing machine learning and deep learning approaches for computational biology.

**Valentina Boeva** is a professor at ETH Zurich, Computer Science Department, Institute for Machine Learning, Zurich, Switzerland. Her research focuses on cancer epigenetics, computational biology, and machine learning.

## References

John P Klein, Melvin L Moeschberger, et al. Survival analysis: techniques for censored and truncated data, volume 1230. Springer, 2003.

Pauline Depuydt, Valentina Boeva, Toby D Hocking, Robrecht Cannoodt, Inge M Ambros, Peter F Ambros, Shahab Asgharzadeh, Edward F Attiyeh, Valérie Combaret, Raffaella Defferrari, et al. Genomic amplifications and distal 6q loss: novel markers for poor survival in high-risk neuroblastoma patients. JNCI: Journal of the National Cancer Institute, 110(10):1084–1093, 2018.

Håvard Kvamme, Ørnulf Borgan, and Ida Scheel. Time-to-event prediction with neural networks and cox regression. arXiv preprint 1907.00825, 2019.

Qixian Zhong, Jonas W Mueller, and Jane-Ling Wang. Deep extended hazard models for survival analysis. In M. Ranzato, A. Beygelzimer, Y. Dauphin, P.S. Liang, and J. Wortman Vaughan, editors, Advances in Neural Information Processing Systems, volume 34, pages 15111–15124. Curran Associates, Inc., 2021. URL https://proceedings.neurips.cc/paper/2021/file/7f6caf1f0ba788cd7953d817724c2b6e-Paper.pdf.

Weijing Tang, Kevin He, Gongjun Xu, and Ji Zhu. Survival analysis via ordinary differential equations. Journal of the American Statistical Association, pages 1–16, 2022.

Changhee Lee, William Zame, Jinsung Yoon, and Mihaela Van Der Schaar. Deephit: A deep learning approach to survival analysis with competing risks. In Proceedings of the AAAI conference on artificial intelligence, volume 32, 2018.

Katarzyna Tomczak, Patrycja Czerwińska, and Maciej Wiznerowicz. The cancer genome atlas (tcga): an immeasurable source of knowledge. Contemporary oncology, 19(1A):A68, 2015.

International Cancer Genome Consortium et al. International network of cancer genome projects. Nature, 464(7291):993, 2010.

Xiaotu Ma, YU Liu, Yanling Liu, Ludmil B Alexandrov, Michael N Edmonson, Charles Gawad, Xin Zhou, Yongjin Li, Michael C Rusch, John Easton, et al. Pan-cancer genome and transcriptome analyses of 1,699 paediatric leukaemias and solid tumours. Nature, 555(7696):371–376, 2018.

Roman Hornung and Marvin N Wright. Block forests: random forests for blocks of clinical and omics covariate data. BMC bioinformatics, 20(1):1–17, 2019.

Moritz Herrmann, Philipp Probst, Roman Hornung, Vindi Jurinovic, and Anne-Laure Boulesteix. Large-scale benchmark study of survival prediction methods using multi-omics data. Briefings in bioinformatics, 22(3):bbaa167, 2021.

Luís A Vale-Silva and Karl Rohr. Long-term cancer survival prediction using multimodal deep learning. Scientific Reports, 11(1):1–12, 2021.

David Wissel, Daniel Rowson, and Valentina Boeva. Systematic comparison of multi-omics survival models reveals a widespread lack of noise resistance. Cell Reports Methods, 3 (4), 2023.

Anika Cheerla and Olivier Gevaert. Deep learning with multimodal representation for pancancer prognosis prediction. Bioinformatics, 35(14):i446–i454, 2019.

Qing Zhao, Xingjie Shi, Yang Xie, Jian Huang, BenChang Shia, and Shuangge Ma. Combining multidimensional genomic measurements for predicting cancer prognosis: observations from tcga. Briefings in bioinformatics, 16(2):291–303, 2015.

Hege M Bøvelstad, Ståle Nygård, and Ørnulf Borgan. Survival prediction from clinico-genomic models-a comparative study. BMC bioinformatics, 10(1):1–9, 2009.

Ziling Fan, Zhangqi Jiang, Hengyu Liang, and Chao Han. Pancancer survival prediction using a deep learning architecture with multimodal representation and integration. Bioinformatics Advances, 3(1):vbad006, 2023.

Roman Hornung, Frederik Ludwigs, Jonas Hagenberg, and Anne-Laure Boulesteix. Prediction approaches for partly missing multi-omics covariate data: A literature review and an empirical comparison study. Wiley Interdisciplinary Reviews: Computational Statistics, page e1626, 2023.

Christina Nießl, Moritz Herrmann, Chiara Wiedemann, Giuseppe Casalicchio, and Anne-Laure Boulesteix. Over-optimism in benchmark studies and the multiplicity of design and analysis options when interpreting their results. Wiley Interdisciplinary Reviews: Data Mining and Knowledge Discovery, 12(2):e1441, 2022.

Sayash Kapoor and Arvind Narayanan. Leakage and the reproducibility crisis in machine-learning-based science. Patterns, 4(9), 2023.

Simon Klau, Vindi Jurinovic, Roman Hornung, Tobias Herold, and Anne-Laure Boulesteix. Priority-lasso: a simple hierarchical approach to the prediction of clinical outcome using multi-omics data. BMC bioinformatics, 19(1):1–14, 2018.

Yi-Kuan Tseng and Ken-Ning Shu. Efficient estimation for a semiparametric extended hazards model. Communications in Statistics—Simulation and Computation®, 40(2):258–273, 2011.

Travers Ching, Xun Zhu, and Lana X Garmire. Cox-nnet: an artificial neural network method for prognosis prediction of high-throughput omics data. PLoS computational biology, 14(4):e1006076, 2018.

Jared L Katzman, Uri Shaham, Alexander Cloninger, Jonathan Bates, Tingting Jiang, and Yuval Kluger. Deepsurv: personalized treatment recommender system using a cox proportional hazards deep neural network. BMC medical research methodology, 18(1):1–12, 2018.

Ethan Cerami, Jianjiong Gao, Ugur Dogrusoz, Benjamin E Gross, Selcuk Onur Sumer, Bülent Arman Aksoy, Anders Jacobsen, Caitlin J Byrne, Michael L Heuer, Erik Larsson, et al. The cbio cancer genomics portal: an open platform for exploring multidimensional cancer genomics data, 2012.

John N Weinstein, Eric A Collisson, Gordon B Mills, Kenna R Shaw, Brad A Ozenberger, Kyle Ellrott, Ilya Shmulevich, Chris Sander, and Joshua M Stuart. The cancer genome atlas pan-cancer analysis project. Nature genetics, 45(10): 1113–1120, 2013.

Jianfang Liu, Tara Lichtenberg, Katherine A Hoadley, Laila M Poisson, Alexander J Lazar, Andrew D Cherniack, Albert J Kovatich, Christopher C Benz, Douglas A Levine, Adrian V Lee, et al. An integrated tcga pan-cancer clinical data resource to drive high-quality survival outcome analytics. Cell, 173(2): 400–416, 2018.

Stef Van Buuren. Flexible imputation of missing data. CRC press, 2018.

Roman Hornung, Christoph Bernau, Caroline Truntzer, Rory Wilson, Thomas Stadler, and Anne-Laure Boulesteix. A measure of the impact of cv incompleteness on prediction error estimation with application to pca and normalization. BMC Medical Research Methodology, 15(1):1–15, 2015.

Laura Antolini, Patrizia Boracchi, and Elia Biganzoli. A time-dependent discrimination index for survival data. Statistics in medicine, 24(24):3927–3944, 2005.

Erika Graf, Claudia Schmoor, Willi Sauerbrei, and Martin Schumacher. Assessment and comparison of prognostic classification schemes for survival data. Statistics in medicine, 18(17-18):2529–2545, 1999.

Humza Haider, Bret Hoehn, Sarah Davis, and Russell Greiner. Effective ways to build and evaluate individual survival distributions. J. Mach. Learn. Res., 21(85):1–63, 2020.

Edward L Kaplan and Paul Meier. Nonparametric estimation from incomplete observations. Journal of the American statistical association, 53(282):457–481, 1958.

Insuk Sohn, Jinseog Kim, Sin-Ho Jung, and Changyi Park. Gradient lasso for cox proportional hazards model. Bioinformatics, 25(14):1775–1781, 2009.

Meritxell Arenas, Sebastià Sabater Marina Gascón, Ivan Henríquez, M José Bueno Àngels Rius, Àngels Rovirosa, David Gómez, Anna Lafuerza, Albert Biete, et al. Quality assurance in radiotherapy: analysis of the causes of not starting or early radiotherapy withdrawal. Radiation Oncology, 9(1):1–6, 2014.

Curtis Northcutt, Anish Athalye, and Jonas Mueller. Pervasive label errors in test sets destabilize machine learning benchmarks. In J. Vanschoren and S. Yeung, editors, Proceedings of the Neural Information Processing Systems Track on Datasets and Benchmarks, volume 1. Curran, 2021. URL https://datasets-benchmarks-proceedings.neurips.cc/paper_files/paper/2021/file/f2217062e9a397a1dca429e7d70bc6ca-Paper-round1.pdf.

