## Supplementary Document for "SurvBoard: standardized benchmarking for multi-omics cancer survival models"

### Supplementary Methods

**A note on standardisation.** We note that our focus on standardisation is meant to apply mainly to the design of our benchmark and less to the results we provide below. Concretely, since model training always requires certain choices (*e.g.,* which hyperparameters the search space should include and what ranges), we believe this part of a benchmark cannot be perfectly standardised. We thus provide what we believe are reasonable default experiments and look forward to other researchers contributing their models (with potentially more diverse modeling approaches) to SurvBoard, while benefiting from our standardised preprocessing and benchmark design.

**Breadth of considered models.** We note that our experiments serve two primary goals: Seed SurvBoard through an initial curated set of models that should give a reasonable approximation to the current state-of-the-art. Second and most importantly, showcasing the benefits of SurvBoard relative to prior work, potentially revealing new findings now and in the future. We would like to emphasize that our focus is on providing the best possible preprocessing and benchmark design, such that the community may provide their predictions to SurvBoard in the future and thus keep track of state of the art performance.

### Datasets

#### The Cancer Genome Atlas (TCGA)

**Background.** The Cancer Genome Atlas (TCGA) [Tomczak et al., 2015] is one of the most impactful large-scale cancer projects in recent years. In addition to clinical data, TCGA provides various omics modalities from 33 cancer types and over 10000 patients, as well as whole slide images. TCGA has also had a large impact on the survival analysis community, with various papers focusing on survival prediction from both multi-omics [Hornung and Wright, 2019, Cheerla and Gevaert, 2019, Vale-Silva and Rohr, 2021] and gene expression [Ching et al., 2018, Kim et al., 2020] data.

**Chosen modalities.** In our work, we used TCGA for survival analysis with multi-omics data, in combination with clinical data. Since our goal was to benchmark the ability of different models to integrate high-dimensional multi-omics data, we chose the highest number of modalities for each dataset in TCGA, while still respecting our other selection criteria. Table S1 gives a full overview of all datasets in TCGA. We did not include whole slide image data in our benchmark, since it is non-trivial to integrate into statistical models, hindering comparisons between neural and statistical methods. Otherwise, we included all available modalities for a particular dataset in TCGA, including gene expression, protein expression, microRNA expression, mutation, DNA methylation, copy number variations, and clinical data.

**Clinical data.** To enable comparability between the standard, missing modality, and the pan-cancer settings, we included the same clinical variables for all datasets on TCGA: Age at diagnosis, gender, race, tumor stage, clinical stage, and histological type. We note that we also included variables that were constant for a particular dataset (*e.g.,* we included gender for breast cancer) to more easily create a pan-cancer dataset. All categories indicating that a particular clinical variable was not recorded or not applicable (or similar) for a particular patient were set to NA and handled accordingly (see Preprocessing).

**Gene expression.** Pan-cancer normalised gene expression data was obtained from PANCANATLAS [Weinstein et al., 2013]. Subsequently, we applied our standard preprocessing (see Preprocessing). We further log transformed gene expression using $\text{expression} = \log_2(\text{expression} + 1)$.

**Protein expression.** Protein expression data was obtained from PANCANATLAS - aside from our standard preprocessing which was applied to all modalities, protein expression was not modified further.

**DNA methylation.** We used *Merged 27K+450K Only* DNA methylation data from PANCANATLAS. No further modality-specific preprocessing was applied.

**Mutation.** We used the *MAF* file available from PANCANATLAS, which we read and transformed into a *gene* $\times$ *patient* matrix using the *mutCountMatrix* function available in *maftools* to calculate the number of non-silent mutations per gene per patient [Mayakonda et al., 2018]. Genes that were non-mutated in a particular dataset were still included for uniformity in the pan-cancer setting. No further modality-specific preprocessing was applied to the mutation data.

**Copy number variations.** For copy number variations, we used the pan-cancer *GISTIC2* [Mermel et al., 2011] thresholded file available on *Xenabrowser* [Goldman et al., 2020] which maps copy number variations to $\{-2, -1, 0, 1, 2\}$. Copy number variations were used without further modality-specific preprocessing.

**MicroRNA.** MicroRNA data were obtained from PANCANATLAS. In addition to our standard preprocessing, we also applied a log transformation to the microRNA expression using $\text{expression} = \log_2(\text{expression} + 1)$.

**Miscellaneous.** If a particular patient on TCGA had multiple (primary) samples available for a specific modality (as may happen, *e.g.,* when multiple vials were used in sequencing), we used the lexicographically smaller vial (*i.e.,* if both vial A and vial B were available, we chose vial A) based on the sample barcode. If there were still multiple primary samples available for that patient, we followed the recommended sort replicate filter by the Broad Institute, choosing the sample with the highest lexicographical sort value (which chose the sample with a higher portion or plate number). Details available at:
https://gdac.broadinstitute.org/runs/stddata__2014_01_15/samples_report/READ_Replicate_Samples.html.

#### International Cancer Genome Consortium (ICGC)

The International Cancer Genome Consortium (ICGC) [Consortium et al., 2010] is a consortium focused on leading various large-scale cancer projects, such as TCGA. Of note, TCGA is a part of ICGC. Within our work, ICGC only refers to the non-TCGA datasets that were available from ICGC. Concretely, we included four datasets from ICGC, namely a lymphotic leukemia dataset from Spain (CLLE-ES), two pancreatic cancer datasets, one from California and from Australia (PACA-CA and PACA-AU) and a liver cancer dataset from Japan (LIRI-JP). All data for our ICGC datasets was obtained from the ICGC data portal. Table S1 provides an overview of all ICGC datasets we included in SurvBoard.

**Chosen modalities.** Based on our general requirements for all datasets, we only included clinical data, gene expression, and mutation for all of our ICGC datasets. We note that while the ICGC data portal includes copy number variation data for our datasets, the segment means (average log2 ratio of probes in that particular segment) were not available for any of our datasets, leading us to exclude copy numbers.

**Clinical data.** We included the following clinical variables for all of our datasets in ICGC (if available): Gender, age at diagnosis, tumor stage, cancer history among family members of the patient, as well as indicators for alcohol and smoking history. Although ICGC did not have a pan-cancer setting, we did not particularly find many differing clinical variables between our datasets and thus restricted ourselves to this set of common clinical variables. All categories indicating that a particular clinical variable was not recorded or not applicable (or similar) for a particular patient were set to NA.

**Gene expression.** For PACA-AU, we used array-based gene expression data, since sequencing-based gene expression was available for only less than 100 patients. For all other datasets, we used sequencing-based gene expression. Sequencing-based gene expression was log-transformed using expression $= \log_2(\text{expression} + 1)$ while array-based gene expression was left untransformed. Aside from this, only our general preprocessing was performed on all gene expression data.

**Mutation.** Mutation data was transformed to *MAF* format using the *icgcSimpleMutationToMAF* function from *maftools*. Afterward, mutation data was transformed into a gene by a patient matrix where for each gene of each patient, we counted the number of non-silent mutations using *mutCountMatrix* from *maftools*. Genes without any mutations for a particular dataset were kept. Otherwise, no modality-specific preprocessing was performed for the mutation data of ICGC.

If a particular patient on ICGC had multiple (primary) samples or specimens available for a specific modality, we chose the specimen that was closest to the time of diagnosis and the sample that had been stored for the shortest amount of time. If two specimens were taken at identical times or two samples had been stored for the same amount of time or no information on sample/specimen time was available, we pseudo-randomly (but seeded) chose one of the available samples/specimens.

### Therapeutically Applicable Research To Generate Effective Treatments (TARGET)

The Therapeutically Applicable Research to Generate Effective Treatments (TARGET) Initiative [GenomeOC, 2021] is a study on childhood cancers that seeks to understand the molecular basis of pediatric malignancy through comprehensive molecular characterisation. The purpose of the study is to accelerate research in novel marker discovery and drug development. The TARGET project currently hosts omics information on 9 pediatric cancers. Following the selection criteria described in the preprocessing section, we ended up with 2 out of the 9 cancers in our benchmark study, namely, Acute Lymphoblastic Leukemia (ALL) Expansion Phase 2, and Wilms Tumor (WT). All datasets are publicly available and can be accessed at `https://portal.gdc.cancer.gov/projects`. The datasets for this benchmark study, however, were accessed through cBioPortal [Cerami et al., 2012, Gao et al., 2013].

**Chosen modalities.** Based on our general requirements for all datasets (see the preprocessing section), we included clinical data, gene expression, copy number variations, and microRNA expression for both TARGET-ALL and clinical data, gene expression, copy number variations, microRNA and DNA methylation for TARGET-WT.

**Clinical data.** Clinical variables were selected such that there was no potential leakage of survival or genetic information. Based on these criteria, we included age at diagnosis, gender, ethnicity, and race for TARGET-ALL and age at diagnosis, clinical stage, gender, ethnicity, and race for TARGET-WT. All categories indicating that a particular clinical variable was not recorded or not applicable (or similar) for a particular patient were grouped into a single "NA" category (except for gender, since having gender not available was exceedingly rare, thus we excluded the few patients that this applied to on TARGET-WT).

**Gene expression.** For all datasets in TARGET, we used sequencing-based RPKM (Reads Per Kilobase of transcript per Million mapped reads) normalised gene expression values of 26, 136 genes. The data was preprocessed as detailed in the main preprocessing section and finally log-transformed using expression $= \log_2(\text{expression} + 1)$.

**Copy number variations.** Putative copy-number calls were determined using *GISTIC2* [Mermel et al., 2011] for all datasets. The mapping is as follows: -2 = homozygous deletion; -1 = hemizygous deletion; 0 = neutral / no change; 1 = gain; 2 = high-level amplification. Copy number variations were directly used without any further modality-specific preprocessing.

**microRNA expression.** The microRNA expression values accessed through cBioPortal for TARGET-ALL and TARGET-WT datasets were preprocessed as detailed in the preprocessing section and finally log-transformed using expression $= \log_2(\text{expression}+1)$.

**DNA methylation.** For the TARGET-WT dataset, methylation beta-values from the HM450 platform were used. The DNA methylation values were used without any further modality-specific preprocessing.

### Molecular Taxonomy of Breast Cancer International Consortium (METABRIC)

The Molecular Taxonomy of Breast Cancer International Consortium (METABRIC) [Curtis et al., 2012] is a study on breast cancer with the aim of defining new subtypes. Molecular Taxonomy of Breast Cancer International Consortium (METABRIC) only consists of one dataset, which also matches our selection criteria. We accessed the data for METABRIC through cBioPortal Cerami et al. [2012], Gao et al. [2013].

**Chosen modalities.** Based on our general requirements for all datasets (see the preprocessing section), we included clinical data, gene expression, mutation, copy number variation, and DNA methylation for METABRIC.

**Clinical data.** Clinical variables were selected such that there was no potential leakage of survival or genetic information. We thus included age at diagnosis, cellularity, menopausal state, laterality, and histology subtype for clinical variables. All categories indicating that a particular clinical variable was not recorded or not applicable (or similar) for a particular patient were grouped into a single "NA" category (if applicable).

**Gene expression.** For METABRIC, we used microarray-based gene expression data. The data was preprocessed as detailed in the main preprocessing section and finally log-transformed using expression $= \log_2(\text{expression} + 1)$ after median imputation.

**Copy number variations.** Putative discrete copy-number calls were used as provided by cBioPortal. No further modality-specific preprocessing was performed for CNVs.

**DNA methylation.** We used DNA methylation values as provided by cBioPortal, which provides "average methylation (beta-values) of promoters defined as 500bp upstream & 50 downstream of Transcription Start Site (TSS)" [Cerami et al., 2012]. No further modality-specific preprocessing was performed for DNA methylation.

**Mutation.** For mutation calls, we used the MAF file provided by cBioPortal and calculated the number of non-silent mutations per gene for each patient using the `mutCountMatrix` function of `maftools` [Mayakonda et al., 2018]. No further modality-specific preprocessing was performed for mutation calls.

### Performance metrics

For a particular patient $i$, let $D_i$ denote their true event time, let $C_i$ denote their right-censoring time, and denote their observed time $T_i = \min(D_i, C_i)$. We denote the censoring indicator as $\delta_i = \mathbb{1}\{T_i = D_i\}$ for the same patient $i$. Lastly, let $\hat{S}(t \mid X_i)$ be the predicted survival time for patient $i$ at time $t$.

Below, we briefly detail the three performance metrics used throughout SurvBoard. We include one metric for discriminative performance, namely Antolini's concordance [Antolini et al., 2005] and two metrics measuring calibration, the Integrated Brier Score [Graf et al., 1999] and D-Calibration [Haider et al., 2020]. Technically, the IBS also (partially) measures discrimination. For a broader discussion on how discriminative and calibration measures differ, as well as the different flavors of calibration metrics, we refer the reader to Chapter 2 of Haider et al. [2020].

**Antolini concordance.** In survival analysis, a pair of samples is said to be concordant if the risk score of the event predicted by the model is lower for the sample who experienced the event at a later time, provided that the pair of samples is comparable. Various concordances may be defined which generally correspond to a (possibly weighted) ratio of all concordance pairs over all comparable pairs of subjects. Concordances thus fall into the $[0, 1]$ interval, where 1 corresponds to a perfect ordering, and 0.5 corresponds to a random ordering, or, equivalently, an ordering that does not differentiate between patients (such as a Kaplan-Meier).

While many recent works in multi-omics survival analysis (*e.g.,* (*e.g.,*) [Herrmann et al., 2021, Hornung and Wright, 2019, Cheerla and Gevaert, 2019]) rely on Harrell's Harrell et al. [1982] or Uno's concordance Uno et al. [2011], we evaluated Antolini's concordance [Antolini et al., 2005], a time-dependent concordance measure (Eq. (1)). Although Antolini's concordance exactly corresponds to Harrell's concordance for proportional-hazards models, it is not the case for other models such as Random Survival Forest (RSF) (see also Sonabend et al. [2022] for a discussion on why a comparison between different concordances should be avoided). Thus, we evaluated Antolini's concordance, primarily to enable SurvBoard to generalize to arbitrary models - models using non-proportional hazards formulations are generally unable to produce per-patient risk measures, which makes evaluating them using Harrell's or Uno's concordance impossible.

$$C_{\text{Antolini}} = \frac{\sum_{i=1}^{n} \sum_{j=1}^{n} \delta_i \mathbb{1}\{\hat{S}(T_i \mid X_i) < \hat{S}(T_i \mid X_j)\}}{\sum_{i=1}^{n} \sum_{j=1}^{n} \delta_i \mathbb{1}\{T_i < T_j\}} \tag{1}$$

Here, $n$ denotes the total number of patients within a particular dataset.

We implemented Antolini's concordance using the default method provided by the *pycox* paper [Kvamme et al., 2019]. The default *pycox* implementation is termed the *adjusted* Antolini concordance and differs very slightly from the original paper [Kvamme et al., 2019, Antolini et al., 2005].

**Integrated Brier Score** The Brier Score (BS), originally conceived for classification [Brier et al., 1950], was extended to right-censored survival data by Graf et al. [1999] (Eq. (2)). Since then, the integrated version, the Integrated Brier score (IBS) (Eq. (3)) has become widely used for measuring the quality of survival predictions, both in genomic applications [Herrmann et al., 2021, Klau et al., 2018] and for low-dimensional clinical survival models [Zhong et al., 2021, Kvamme et al., 2019]. A lower Integrated Brier score (IBS) indicates better model performance.

$$\begin{aligned} BS(t) = \frac{1}{n} \sum_{i=1} & \frac{\hat{S}(t \mid X_i)^2 \mathbb{1}\{T_i \leq t \cap \delta_i = 1\}}{\hat{C}(T_i)} \\ & + \frac{(1 - \hat{S}(t \mid X_i))^2 \mathbb{1}\{T_i > t\}}{\hat{C}(t)} \end{aligned} \tag{2}$$

Here, $\hat{G}(t \mid X_i)$ is an estimate of the censoring survival function, *i.e.,* (*i.e.,*) $\hat{G}(t \mid X_i) = P(C_i > t \mid X_i)$ [Kvamme et al., 2019].

$$IBS = \frac{1}{b - a} \int_a^b BS(x)dx \tag{3}$$

We used the *pycox* implementation of IBS. In particular, we set $a$ equal to the minimum of all test times in each split and $b$ to the maximum of all test times in each split and considered a time grid of 100 points between the minimum and maximum time to approximate the true integral. Further, we set $\hat{G}(t \mid X_i) = \hat{G}(t)$ to be the Kaplan-Meier estimator on the test set.

**D-Calibration (D-CAL)** D-Calibration (D-CAL) is a rather new metric focused on evaluating the distributional calibration of *individual survival curves*. We refer the reader to Haider et al. [2020] for the full algorithm.

We computed D-Calibration using the *survival_evaluation* package provided by the paper authors [Haider et al., 2020]. Throughout our results, we report test statistics instead of p-values for better comparability, which implies that lower values signify better performance. We used D-CAL with its default parameter of 10 bins. In case this created missing values for a specific split of a particular model, we replaced the missing value with the mean D-CAL across all other splits for the same model.

### SurvBoard walk-through

Here, we provide a short example walk-through on our web service, SurvBoard, by submitting an example file to it. More detailed documentation is available under the documentation tab of the web service (Fig. S1).

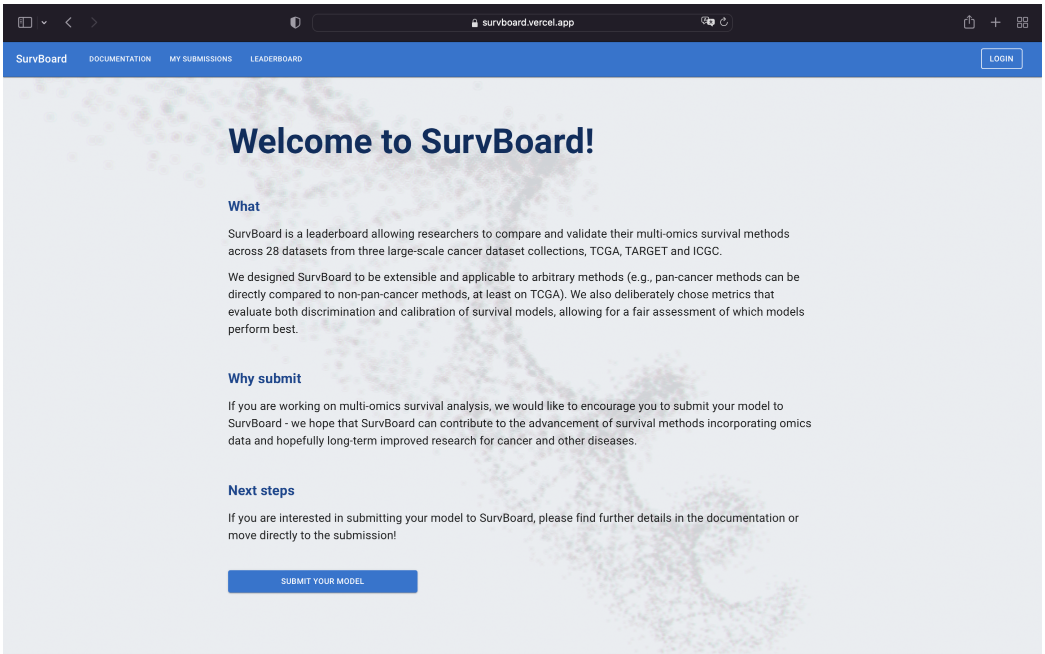


**Fig. S1.** The welcome page of our web service provides further documentation on how to submit and other details.

#### Leaderboard

Our leaderboard was designed with a simple spreadsheet structure in mind - users may easily subset all models to only include a particular setting or datasets from a particular project Fig. S2. To avoid overwhelming users, SurvBoard only displays one metric at a time, showing the mean of a metric across test splits and its standard deviation in parentheses, both rounded to three digits, per dataset. We also included an aggregated column that shows the pooled mean and standard deviation across datasets for a particular project. The public leaderboard is freely accessible, without a need to sign-up.

#### Sign-up

If users want to submit their models to SurvBoard, they are required to sign-up with a name, email, and institution/organisation (Fig. S3). We require these details to communicate with users and ensure authenticity such that repeated submissions to game SurvBoard and the usage of automated systems is not possible.

#### Downloading data and splits

We provide users with the option of downloading all the datasets we used in our benchmark, including the splits for creating the test set. However, signing up is not necessary for simply downloading the data which can instead be accessed directly from Zenodo.

#### Submitting predictions

Once users have trained their respective models and made predictions in our required format (please refer to our web service documentation for details), model predictions may be uploaded through a simple three-step workflow on SurvBoard (Fig. S4 - Fig. S6).

After submission, local results (that is, results only available to the user who submitted them) will generally be available within minutes. We note that we require code submission (see Fig. S5) in the form of an online code repository for submissions that will

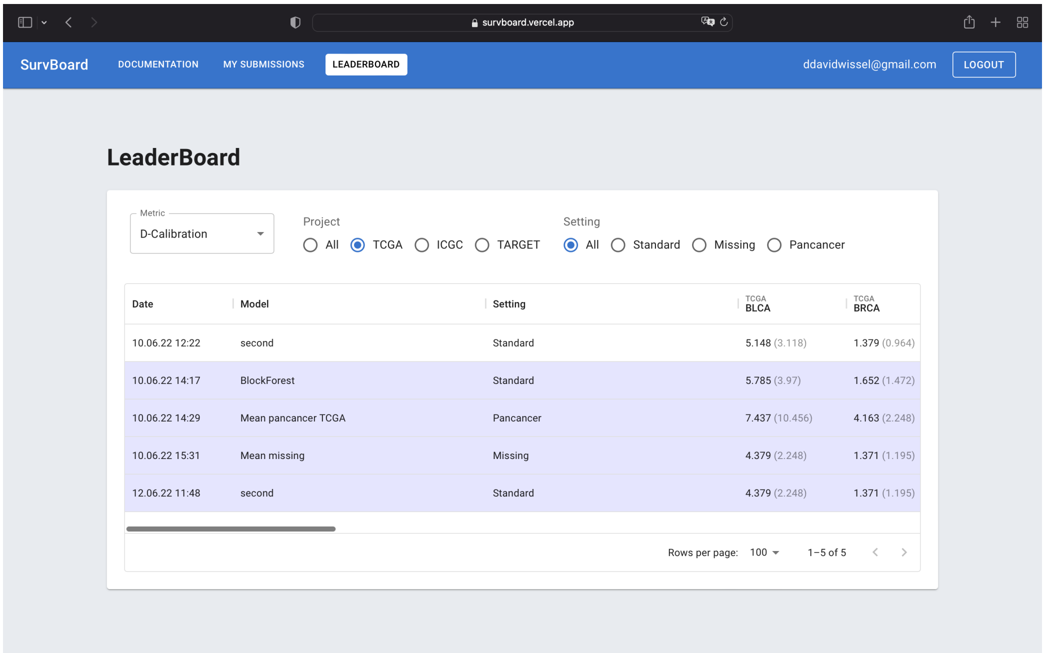


**Fig. S2.** The SurvBoard leaderboard can be easily subset to different projects, settings and display one of three metrics (see Metrics for an overview).

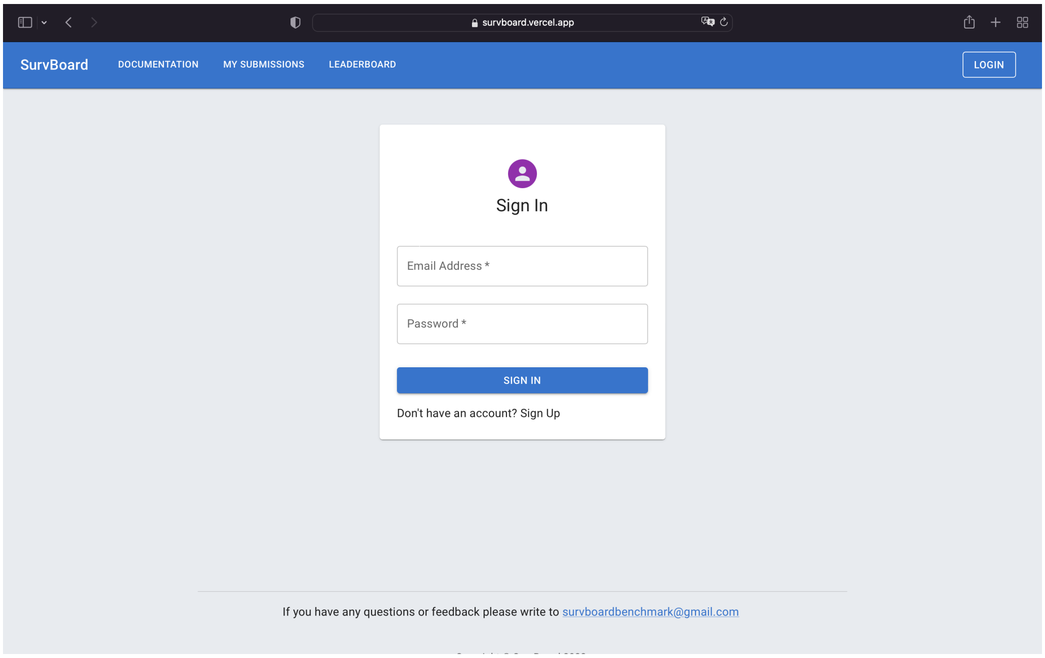


**Fig. S3.** If users want to submit predictions, we require a registration, in order to prevent malicious usage.

be shown publicly. In case users opt for public submission, by submitting a link to a code repository implementing their chosen model, results will be available on the public leaderboard after review for potential overfitting or of previous submission attempts to SurvBoard, which will generally take less than a week.

### Statistical models

**Reproducibility.** Results for all statistical results may be reproduced using requisite scripts within our Github repository. For completeness, we provide information on which hardware each of the statistical models was run in the sections below. All of our models were interfaced using *mlr3* [Lang et al., 2019] and *mlr3proba* [Sonabend et al., 2021].

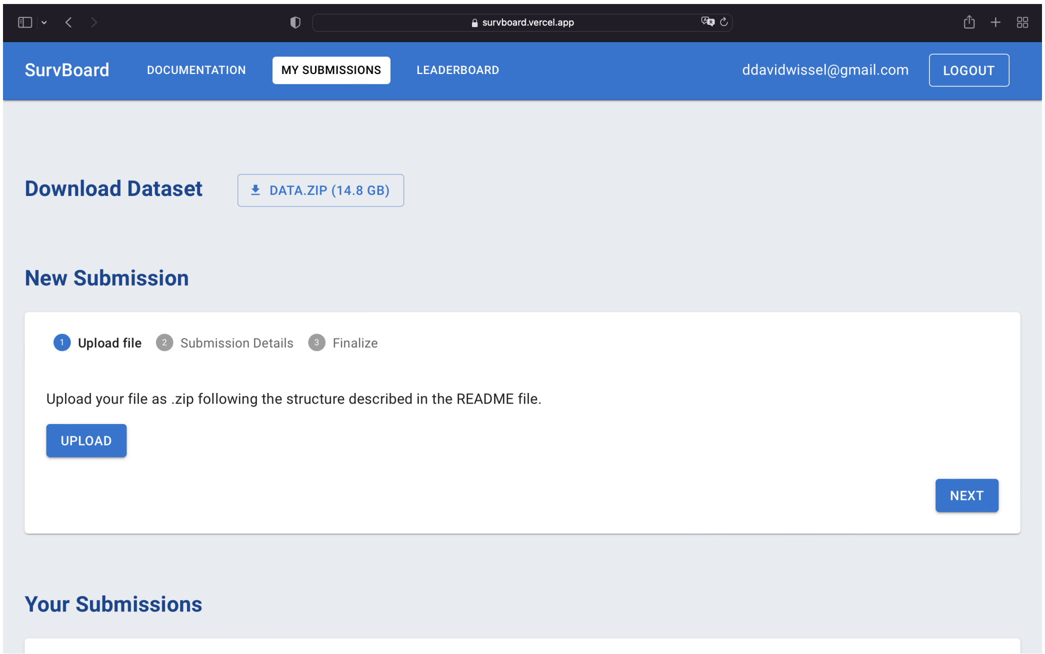


**Fig. S4.** Initially, SurvBoard requires the user to upload a ZIP file containing model predictions for one or more datasets.

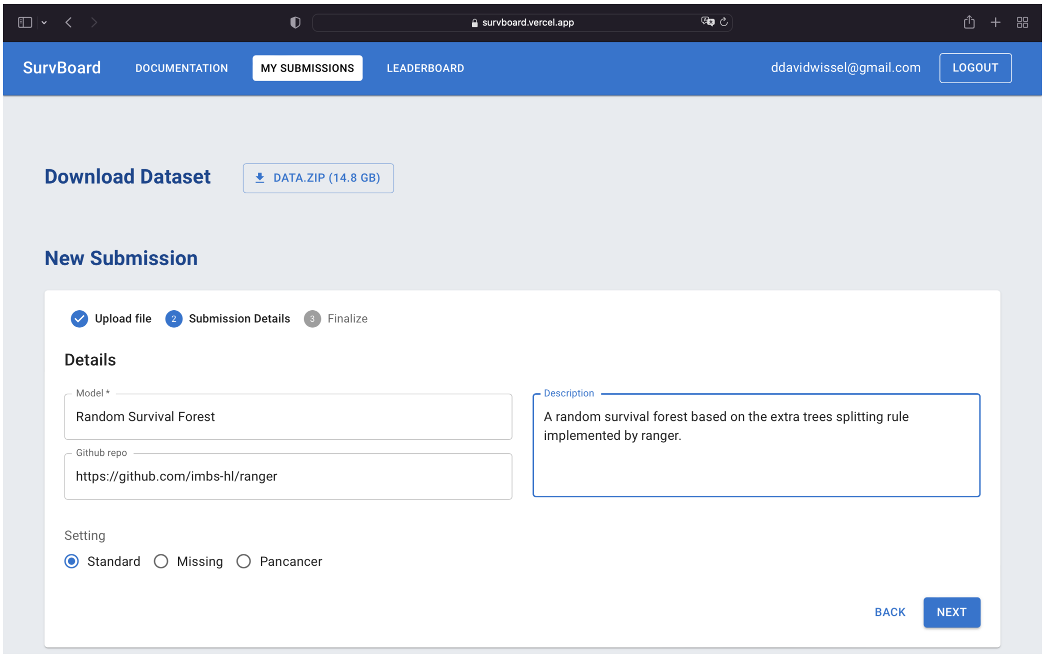


**Fig. S5.** In the second step of submission, the user may give their model a name, and describe what their model is based on. Further, if users want to opt for a public release of their model performance, we also require the submission of a code repository link.

**Licenses.** All considered statistical models are available on CRAN or other repositories as noted in our *renv* lock file [Ushey and Wickham, 2024]. To the best of our knowledge, all statistical models used open-source licenses at the time of our experiments.

**PriorityLasso L1+L2.** We included PriorityLasso L1+L2, a recently proposed method for survival prediction from multi-modal data, in our benchmark primarily due to its good performance in the benchmark of Herrmann et al. [2021]. We use the authors' official R package for implementation [Klau et al., 2018]. Specifically, we used the approach described by Herrmann et al. [2021] to choose the block order, first fitting an $\ell_2$ penalized Cox model on variables that only belong to each of the considered modalities. Afterwards, we chose the first block to be the modality with the highest absolute mean coefficients, the second block to be the modality with the second highest value, and so on. The modality-specific regularization hyperparameters were chosen based on the maximization of partial (log)-likelihood on the validation folds (*i.e.*, the test folds of our *inner* cross-validation, which is used for

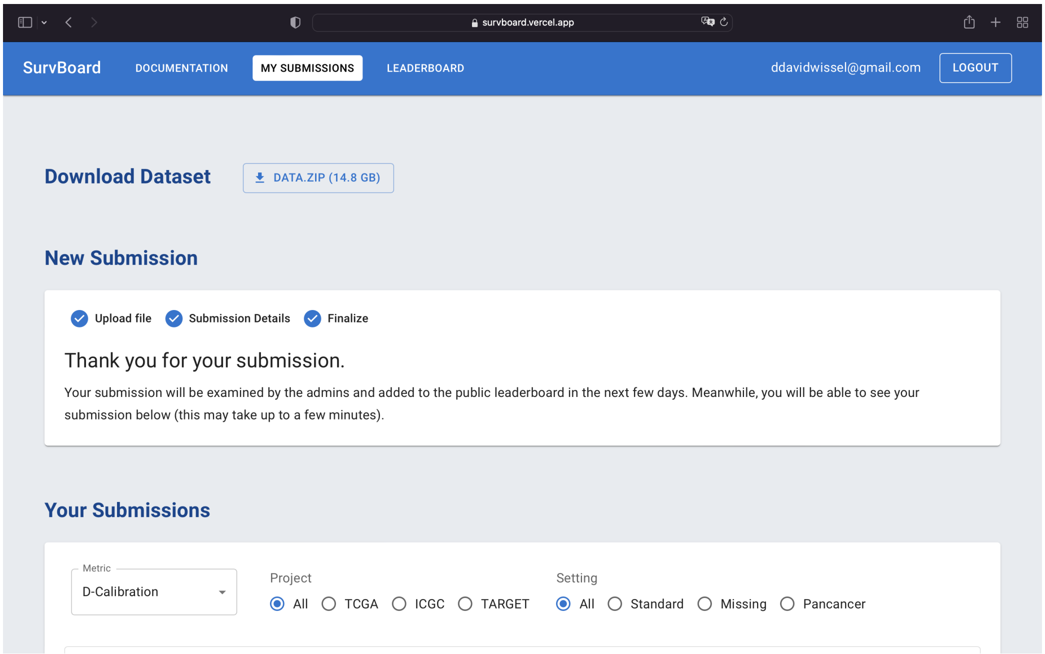


**Fig. S6.** Lastly, users are informed that it may take a few days until their model performance is visible publicly - although they can immediately see their performance on SurvBoard locally.

tuning) (*lambda.type="lambda.min"* and *type.measure="deviance"*). We used five-fold cross-validation, stratified by the event indicator for choosing the hyperparameters for each block. Contrary to Herrmann et al. [2021], we also use five-fold cross-validation to estimate the offsets - (*cvoffset=TRUE* and *cvoffsetnfolds=5*). Lastly, we dummy-encoded categorical variables and standardized all input variables using *PriorityLasso L1+L2* (*i.e., standardize = TRUE*). To prevent model failures sometimes encountered by the Lasso and guard against other failure cases of the Lasso, we ran PriorityLasso L1+L2 using an elastic net with a ridge component of 0.1 (*i.e.,* $\alpha = 0.9$) All other settings of *PriorityLasso L1+L2* were left at their defaults. The search space for the regularization hyperparameter $\lambda$ for both PriorityLasso L1+L2 and the initial step to choose the priority order used the sequence approach in *glmnet* [Friedman et al., 2010, Simon et al., 2011], which chooses 100 values for $\lambda$ between a completely sparse fit (for $\lambda_{\text{max}}$) and a minimum $\lambda_{\text{min}} = 0.01\lambda_{\text{max}}$. All experiments for PriorityLasso L1+L2 were run on a cluster using AMD EPYC-Rome CPUs. PriorityLasso L1+L2 offers native support for running in the missing modality setting for which we use the complete cases approach to impute offsets [Klau et al., 2018, Hornung et al., 2024].

**Block forest.** Since it was among the best-performing models in the study of Herrmann et al. [2021], we also included Block forest in our benchmark, a random forest variant adapted to group structured (*e.g.,* multi-omics) data. We use the authors' official R package for implementation [Hornung and Wright, 2019]. We used the function provided for hyperparameter tuning in the authors' package called *blockfor*. The *Block forest* package tunes hyperparameters using out-of-bag estimates of Harrell's concordance and chose the hyperparameter weight set which maximized the out-of-bag estimate of Harrell's concordance [Harrell et al., 1982]. All other parameters were left at their defaults except that during tuning we only considered 100 trees in each forest (`num.trees.pre = 100`). Categorical variables were passed to Block forest directly and numerical covariates were not standardised. Experiments were run on a cluster using AMD EPYC-Rome CPUs. Since Block forest does not natively account for missing modalities and it is unclear what is the best way to do so, it was not run in the missing modality setting. This is consistent with related work [Hornung et al., 2024].

#### Miscellaneous

We note that for PriorityLasso L1+L2, we used the *Breslow* estimator of the *survival* R package to calculate survival function estimates. Survival functions for Block forest were calculated using package-internal prediction functions which estimate survival functions in each leaf node using a Kaplan-Meier estimator. Copy number was treated as numerical (as opposed to categorical) for all statistical models.

### Deep learning models

**Reproducibility.** We followed best practices in *Pytorch* for reproducibility. Results may be reproduced using requisite scripts within our Github repository. However, we note that hardware differences or multi-threading may cause results to be similar but not exact. For completeness, we provide information on which hardware each of the neural models was run in the sections below.

**Licenses.** All neural models were implemented in *Pytorch* [Paszke et al., 2019] and *skorch* [Tietz et al., 2017] both of which are open source and freely available on Github, via conda, or via pip. We refer to the respective code repositories of *Pytorch* and *skorch* for details on their respective licenses.

**Model details.** All models were fit using batch normalisation [Ioffe and Szegedy, 2015] and ReLUs [Nair and Hinton, 2010] as activation functions. In addition, all models were trained using the AdamW [Loshchilov and Hutter, 2018] optimiser. We used 20% of the training set as a validation set for early stopping (using patience of 10 and restoring the model after the epoch with the lowest validation loss).

Hyperparameters were tuned using five-fold cross-validation stratified by the event indicator and choosing the dropout probability that yielded the lowest negative partial log-likelihood across validation folds. We started out using the hyperparameter grid as Zhong et al. [2021] but had to narrow it for some parameters due to numerical issues that arose due to the high dimensionality and low sample size of our datasets. In particular, we fixed batch size at 1024, the number of hidden layers at one and tuned the learning rate in $\{0.0005, 0.0008, 0.001, 0.005\}$, weight decay in $\{0.0005, 0.005, 0.05, 0.1\}$, the number of hidden nodes in $\{32, 64, 128, 256, 512\}$ and the dropout probability in $\{0, 0.2, 0.4, 0.6\}$.

**Loss functions.**

We assume $T_i$ and $C_i$ to denote the event and right-censoring times of patient $i$, respectively. In right-censored observational survival analysis, we have triplets $(x_i, \delta_i, O_i)$, where $O_i = \min(T_i, C_i)$ and $\delta_i = \mathbb{1}(T_i \leq C_i)$

In addition, here, we assume $\tilde{T}$ is a set of unique, ordered (ascending) event times. Further, $R_i$ is the risk set at time $i$, that is, $R(i) = \{j : O_j \geq O_i\}$. Lastly, $D_i$ denotes the death set at time $i$, $D(i) = \{j : O_j = i \wedge \delta_i = 1\}$.

We wrote torch loss functions for both the Cox PH partial likelihood Eq. (4) and the EH likelihood Eq. (5). For the Cox PH partial likelihood, we used the Breslow approximation to match PriorityLasso L1+L2. For the EH model, we used the standard Gaussian kernel and set the bandwidth to $b = (8\sqrt{2}/3)^{1/5} n^{-1/5}$ as done by Zhong et al. [2021] in their experiments.

$$\ell(h)_{\text{Breslow}} = \sum_{i:i\in\tilde{T}} \sum_{j:j\in D(i)} h_j - |D(j)| \log\left(\sum_{j:j\in R_i} \exp(h_j)\right) \tag{4}$$

$$\begin{aligned} &\ell_{\text{EH}}(h_1, h_2) = \\ &\frac{1}{n}\sum_{i=1}^{n} \delta_i \left[h_2(X_i) - R_i(h_1)\right] \\ &+ \frac{1}{n}\sum_{i=1}^{n} \delta_i \log\left[\frac{1}{nb}\sum_{j=1}^{n} \delta_j K\left(\frac{R_j(h_1) - R_i(h_1)}{b}\right)\right] \\ &- \frac{1}{n}\sum_{i=1}^{n} \delta_i \log\left[\frac{1}{n}\sum_{j=1}^{n} \frac{e^{h_2(X_j)}}{e^{h_1(X_j)}} \Phi\left(\frac{R_j(h_1) - R_i(h_1)}{b}\right)\right] \end{aligned} \tag{5}$$

**Fusion methods.** We used the two best-performing fusion methods established by [Wissel et al., 2023], namely late fusion using the arithmetic mean and intermediate fusion using concatenation to merge the modality-specific representations. We used the implementations provided by Wissel et al. [2023] and refer to their work for notation and implementation details.

**Miscellaneous.** For all neural models, we one-hot encoded categorical variables and standardised all covariates using *StandardScaler* from *scikit-learn* [Pedregosa et al., 2011]. Categories that were present in a particular test split but not the training split were ignored (*i.e.,* all columns for that particular one-hot encoded category were set to zero for unknown categories, using *handle_unknown=ignore* in *OneHotEncoder*). The survival functions for all Cox PH-based models were estimated using the Breslow estimator implementation from *scikit-survival* [Pölsterl, 2020]. For EH-based models, we estimated survival functions via numerical integration of the hazard function using the Gaussian quadrature implementation of *scipy* (`scipy.integrate.quadrature`). For handling missing modalities with all deep learning methods, we used a simple imputation approach. In particular, we set all modalities that were missing in the train to zero *after* preprocessing. This technique is consistent with previous work and has occasionally also been amended with a so-called modality dropout which aims to make models more robust to missing modalities by occasionally dropping out present modalities during training [Cheerla and Gevaert, 2019, Vale-Silva and Rohr, 2021]. We did not use modality dropout here to increase comparability with the non-missing modality setting.

### Maintenance plan

SurvBoard is currently deployed in the cloud on a monthly subscription basis. The service uses common deployment patterns such as containers and SQL datastores and so fits the definition of a hybrid cloud application. The service can be migrated to other vendors if and when needed. The web service deployment fees are currently covered by internal lab funds. Security patches, package updates, and similar tasks will be handled by the paper authors regardless of the deployment environment. The user provides us with an organisational email, which is public knowledge, to be able to create a profile and map their submission correctly. The results from the user's model are used to compute a score and place them on the leaderboard after which the result files are deleted immediately. We only store the user email and the final leaderboard scores for display. Code is only collected via Github repository links, *i.e.,*, all code submitted by users is anyway public on the respective Github repository under the respective license.

The datasets are available on Zenodo: https://zenodo.org/records/11066227. The two co-first authors will share joint responsibility for the web service maintenance for the next three years; after which responsibility will be handled jointly by the appropriate members of one of the two labs.

### Supplementary tables

**Table S1.** Overview of all datasets considered in our benchmark. GEX denotes gene expression, CNV denotes copy number variation, and RPPA denotes protein expression. Modality columns denote the dimensionality of each modality in each specific dataset. $n$ denotes number of samples and $e$ denotes number of events. Complete and incomplete denote the numbers of complete modality and incomplete modality samples, respectively.

| Cancer type | Project | # modalities | # clinical | # GEX | # mutation | # DNA methylation | # CNV | # RPPA | # miRNA | $p$ | $n$ (complete) | $e$ (incomplete) | $n$ (incomplete) | $e$ (incomplete) |
|---|---|---|---|---|---|---|---|---|---|---|---|---|---|---|
| BRCA | METABRIC | 5 | 5 | 20387 | 173 | 11823 | 22542 | 0 | 0 | 54933 | 1376 | 815 | 603 | 328 |
| BLCA | TCGA | 7 | 5 | 20531 | 19687 | 22584 | 24776 | 189 | 743 | 88518 | 330 | 149 | 80 | 31 |
| BRCA | TCGA | 7 | 5 | 20531 | 19687 | 22591 | 24776 | 190 | 743 | 88526 | 771 | 103 | 312 | 48 |
| COAD | TCGA | 7 | 5 | 17507 | 19687 | 22583 | 24776 | 191 | 743 | 85495 | 289 | 64 | 153 | 34 |
| ESCA | TCGA | 7 | 6 | 19076 | 19687 | 22543 | 24776 | 193 | 743 | 87027 | 123 | 44 | 62 | 33 |
| HNSC | TCGA | 7 | 6 | 20531 | 19687 | 22589 | 24776 | 191 | 743 | 88526 | 202 | 122 | 325 | 101 |
| KIRC | TCGA | 7 | 5 | 20531 | 19687 | 22564 | 24776 | 191 | 743 | 88500 | 312 | 88 | 223 | 89 |
| KIRP | TCGA | 7 | 5 | 20531 | 19687 | 22580 | 24776 | 194 | 743 | 88519 | 203 | 29 | 85 | 15 |
| LGG | TCGA | 7 | 5 | 20531 | 19687 | 22588 | 24776 | 190 | 743 | 88523 | 412 | 95 | 99 | 30 |
| LUAD | TCGA | 7 | 5 | 20531 | 19687 | 22587 | 24776 | 189 | 743 | 88521 | 340 | 136 | 169 | 47 |
| PAAD | TCGA | 7 | 6 | 20531 | 19687 | 22537 | 24776 | 190 | 743 | 88473 | 109 | 61 | 75 | 39 |
| SARC | TCGA | 7 | 5 | 20531 | 19687 | 22533 | 24776 | 193 | 743 | 88471 | 194 | 70 | 67 | 29 |
| SKCM | TCGA | 4 | 6 | 20531 | 19687 | 22579 | 0 | 0 | 0 | 62806 | 102 | 29 | 352 | 185 |
| STAD | TCGA | 7 | 5 | 19076 | 19687 | 22577 | 24776 | 193 | 743 | 87060 | 304 | 126 | 113 | 42 |
| UCEC | TCGA | 7 | 5 | 17507 | 19687 | 22598 | 24776 | 191 | 743 | 85510 | 395 | 63 | 151 | 28 |
| OV | TCGA | 7 | 5 | 19076 | 19687 | 22521 | 24776 | 191 | 743 | 87002 | 166 | 98 | 416 | 250 |
| LIHC | TCGA | 7 | 5 | 20531 | 19687 | 22579 | 24776 | 190 | 743 | 88514 | 157 | 80 | 214 | 52 |
| LUSC | TCGA | 7 | 5 | 20531 | 19687 | 22591 | 24776 | 189 | 743 | 88525 | 288 | 117 | 209 | 98 |
| LAML | TCGA | 5 | 6 | 16765 | 0 | 22594 | 24776 | 0 | 743 | 64887 | 139 | 86 | 34 | 22 |
| CESC | TCGA | 7 | 5 | 20531 | 19687 | 22565 | 24776 | 193 | 743 | 88503 | 141 | 24 | 153 | 47 |
| GBM | TCGA | 4 | 6 | 20531 | 19687 | 0 | 24776 | 0 | 0 | 65003 | 144 | 114 | 451 | 377 |
| READ | TCGA | 6 | 6 | 17507 | 19687 | 22582 | 24776 | 0 | 743 | 85304 | 125 | 17 | 39 | 9 |
| LIRI-JP | ICGC | 3 | 4 | 22848 | 6828 | 0 | 0 | 0 | 0 | 29683 | 229 | 42 | 31 | 4 |
| PACA-CA | ICGC | 3 | 6 | 65802 | 6041 | 0 | 0 | 0 | 0 | 71852 | 162 | 128 | 63 | 36 |
| PACA-AU | ICGC | 3 | 4 | 47265 | 5204 | 0 | 0 | 0 | 0 | 52476 | 223 | 134 | 227 | 153 |
| CLLE-ES | ICGC | 3 | 6 | 100090 | 3116 | 0 | 0 | 0 | 0 | 103215 | 294 | 46 | 230 | 34 |
| ALL | TARGET | 4 | 4 | 23716 | 0 | 0 | 20634 | 0 | 1870 | 46227 | 125 | 71 | 1410 | 183 |
| WT | TARGET | 5 | 5 | 22834 | 0 | 15018 | 20634 | 0 | 1870 | 60364 | 116 | 47 | 535 | 67 |

**Table S2.** Tissue sites and full names of all cancer abbreviations used throughout our benchmark.

| Project | Abbreviation | Tissue | Full name |
|---|---|---|---|
| TCGA | BLCA | Bladder | Bladder Urothelial Carcinoma |
| TCGA | BRCA | Breast | Breast invasive carcinoma |
| TCGA | COAD | Colon | Colon adenocarcinoma |
| TCGA | ESCA | Esophagus | Esophageal carcinoma |
| TCGA | HNSC | Head and Neck | Head and Neck squamous cell carcinoma |
| TCGA | KIRC | Kidney | Kidney renal clear cell carcinoma |
| TCGA | KIRP | Kidney | Kidney renal papillary cell carcinoma |
| TCGA | LGG | Brain | Brain Lower Grade Glioma |
| TCGA | LUAD | Lung | Lung adenocarcinoma |
| TCGA | PAAD | Pancreas | Pancreatic adenocarcinoma |
| TCGA | SARC | Bones and soft tissue | Sarcoma |
| TCGA | SKCM | Skin | Skin Cutaneous Melanoma |
| TCGA | STAD | Stomach | Stomach adenocarcinoma |
| TCGA | UCEC | Uterus | Uterine Corpus Endometrial Carcinoma |
| TCGA | OV | Ovaries | Ovarian serous cystadenocarcinoma |
| TCGA | LIHC | Liver | Liver hepatocellular carcinoma |
| TCGA | LUSC | Lung | Lung squamous cell carcinoma |
| TCGA | LAML | Blood forming tissue | Acute Myeloid Leukemia |
| TCGA | CESC | Cervix | Cervical squamous cell carcinoma and endocervical adenocarcinoma |
| TCGA | GBM | Brain and spine | Glioblastoma multiforme |
| TCGA | READ | Rectum | Rectum adenocarcinoma |
| ICGC | CLLE | Blood forming tissue | Chronic Lymphocytic Leukemia |
| ICGC | PACA-AU | Pancreas | Pancreatic Cancer |
| ICGC | PACA-CA | Pancreas | Pancreatic Cancer |
| ICGC | LIRI | Liver | Liver Cancer |
| TARGET | ALL | White blood cells | Acute lymphoblastic leukemia |
| TARGET | WT | Kidney | Wilms tumor |
| METABRIC | BRCA | Breast | Breast invasive carcinoma |

**Table S3.** 'Failure rates' in percentage per model and modalities in our main benchmark results. Combinations of models and modalities that had no failures were not shown. We note that high failure rates for PriorityLasso L1+L2 imply that the best regularization hyperparameter often led to a fully sparse model. In these cases, we fall back to a Kaplan-Meier estimator.

| Model | Modalities | Failure rate |
|---|---|---|
| PriorityLasso L1+L2 | Clinical | 0.153 |
| PriorityLasso L1+L2 | CNV | 0.428 |
| PriorityLasso L1+L2 | GEX | 0.271 |
| PriorityLasso L1+L2 | DNA methylation | 0.362 |
| PriorityLasso L1+L2 | miRNA | 0.290 |
| PriorityLasso L1+L2 | Mutation | 0.478 |
| PriorityLasso L1+L2 | RPPA | 0.294 |
| PriorityLasso L1+L2 | Clinical + GEX | 0.066 |
| PriorityLasso L1+L2 | All modalities | 0.006 |

**Table S4.** Number of failures in D-Calibration calculation per model and modalities in our main benchmark results. Combinations of models and modalities that had no failures were not shown. We note that these NAs were created in an upstream package due to numerical issues for some of the estimated survival functions

| Model | Modalities | # Failures |
|---|---|---|
| NN Cox IC | GEX | 17 |
| NN Cox IC | miRNA | 1 |
| NN Cox IC | Mutation | 5 |
| NN EH IC | GEX | 5 |
| NN EH IC | miRNA | 1 |
| NN EH IC | Mutation | 1 |
| PriorityLasso L1+L2 | Clinical | 1 |
| PriorityLasso L1+L2 | GEX | 5 |
| PriorityLasso L1+L2 | DNA methylation | 5 |
| PriorityLasso L1+L2 | Mutation | 7 |
| NN Cox IC | Clinical + GEX | 18 |
| NN Cox IC | All modalities | 24 |
| NN Cox LM | Clinical + GEX | 19 |
| NN Cox LM | All modalities | 21 |
| NN EH IC | Clinical + GEX | 5 |
| NN EH IC | All modalities | 7 |
| NN EH LM | Clinical + GEX | 9 |
| NN EH LM | All modalities | 8 |
| PriorityLasso L1+L2 | Clinical + GEX | 1 |
| PriorityLasso L1+L2 | All modalities | 7 |
| NN Cox IC | All modalities (missing) | 20 |
| NN Cox LM | All modalities (missing) | 21 |
| NN EH IC | All modalities (missing) | 3 |
| NN EH LM | All modalities (missing) | 6 |
| PriorityLasso L1+L2 | All modalities (missing) | 7 |

### Supplementary figures

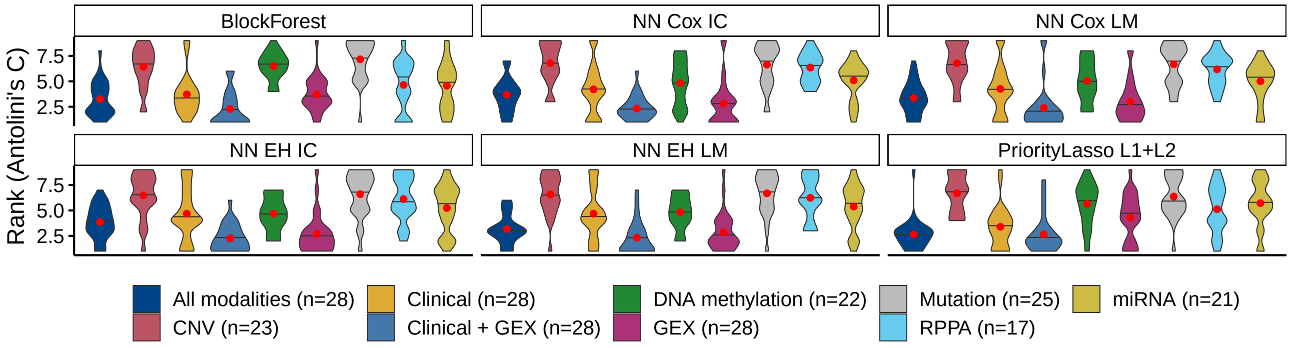


**Fig. S7.** Overall ranked results per model, across modalities in terms of Antolini's concordance (Antolini's C). We note that not all modalities are present in some datasets, (see $n = \ldots$). The median is indicated by black horizontal lines, the arithmetic mean by red points.

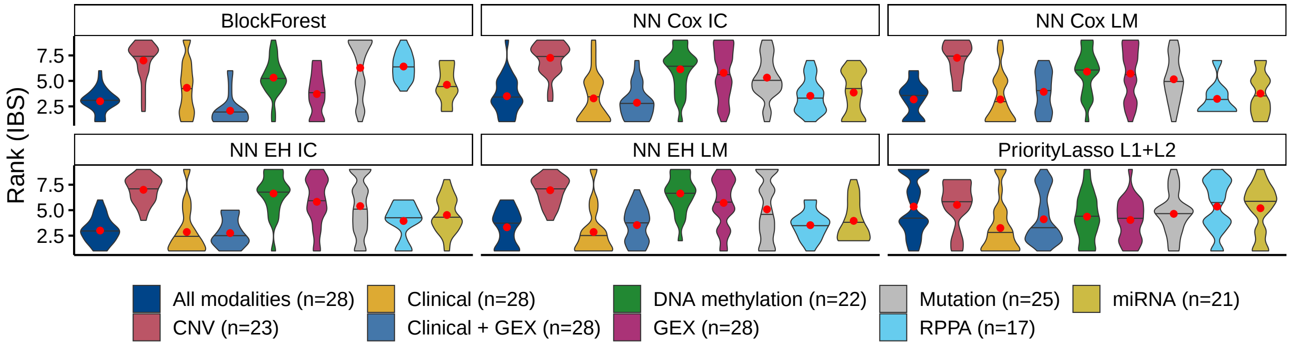


**Fig. S8.** Overall ranked results per model, across modalities in terms of the Integrated Brier Score (IBS). We note that not all modalities are present in some datasets, (see $n = \ldots$). The median is indicated by black horizontal lines, the arithmetic mean by red points.

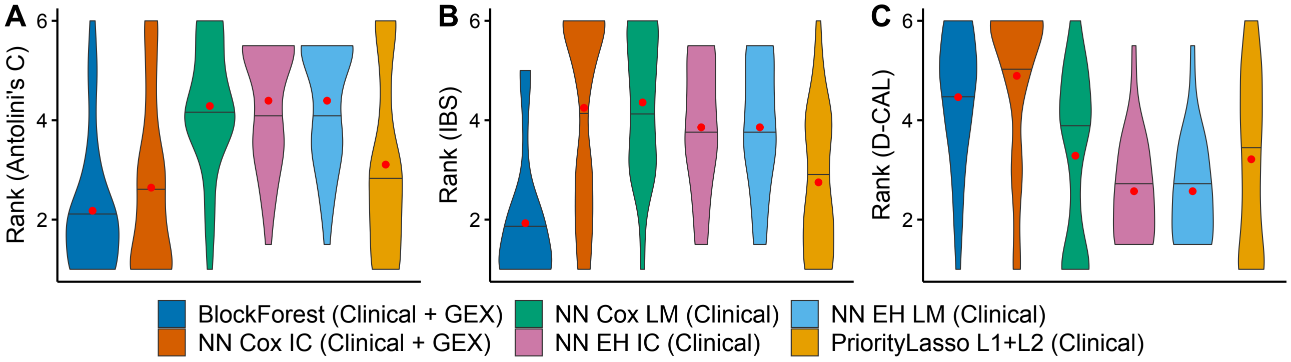


**Fig. S9.** Overall ranked results across all datasets, where lower ranks indicate better scores. A. Antolini's concordance (Antolini's C). B. Integrated Brier Score (IBS). C. D-Calibration (D-CAL). Each model was trained on the dataset on which it ranked the best among itself in terms of the median IBS rank (see Supplementary Fig. S8). Ties in the median IBS rank were broken using the mean IBS rank. The median is indicated by black horizontal lines, the arithmetic mean by red points.

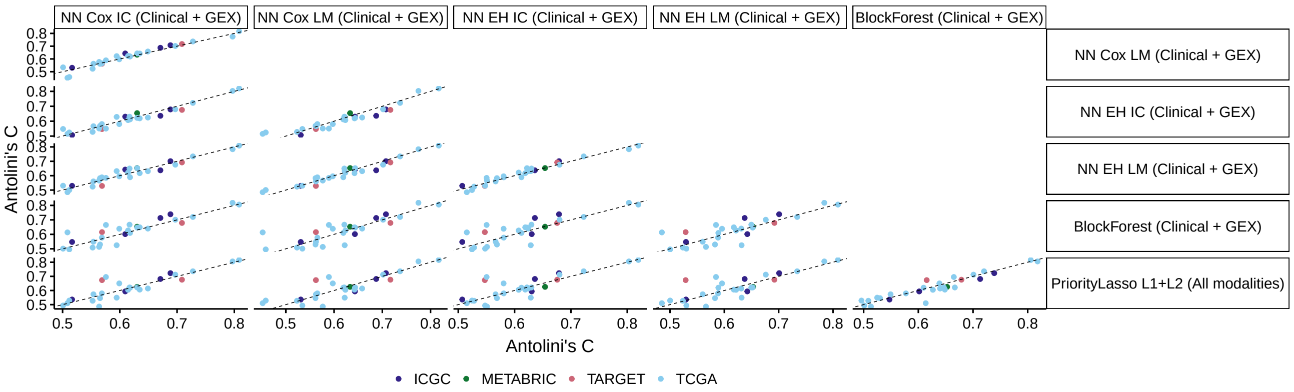


**Fig. S10.** Paired scatter plots of Antolini's C between all methods in terms of Antolini's C. Each model was trained on the modalities on which it ranked the best among itself in terms of Antolini's concordance (see Supplementary Fig. S7).

### References


Katarzyna Tomczak, Patrycja Czerwińska, and Maciej Wiznerowicz. The cancer genome atlas (tcga): an immeasurable source of knowledge. *Contemporary oncology*, 19(1A):A68, 2015.

Roman Hornung and Marvin N Wright. Block forests: random forests for blocks of clinical and omics covariate data. *BMC bioinformatics*, 20(1):1–17, 2019.

Anika Cheerla and Olivier Gevaert. Deep learning with multimodal representation for pancancer prognosis prediction. *Bioinformatics*, 35(14):i446–i454, 2019.

Luís A Vale-Silva and Karl Rohr. Long-term cancer survival prediction using multimodal deep learning. *Scientific Reports*, 11(1): 1–12, 2021.

Travers Ching, Xun Zhu, and Lana X Garmire. Cox-nnet: an artificial neural network method for prognosis prediction of high-throughput omics data. *PLoS computational biology*, 14(4):e1006076, 2018.

Sunkyu Kim, Keonwoo Kim, Junseok Choe, Inggeol Lee, and Jaewoo Kang. Improved survival analysis by learning shared genomic information from pan-cancer data. *Bioinformatics*, 36(Supplement_1):i389–i398, 2020.

John N Weinstein, Eric A Collisson, Gordon B Mills, Kenna R Shaw, Brad A Ozenberger, Kyle Ellrott, Ilya Shmulevich, Chris Sander, and Joshua M Stuart. The cancer genome atlas pan-cancer analysis project. *Nature genetics*, 45(10):1113–1120, 2013.

Anand Mayakonda, De-Chen Lin, Yassen Assenov, Christoph Plass, and H Phillip Koeffler. Maftools: efficient and comprehensive analysis of somatic variants in cancer. *Genome research*, 28(11):1747–1756, 2018.

Craig H Mermel, Steven E Schumacher, Barbara Hill, Matthew L Meyerson, Rameen Beroukhim, and Gad Getz. Gistic2. 0 facilitates sensitive and confident localization of the targets of focal somatic copy-number alteration in human cancers. *Genome biology*, 12 (4):1–14, 2011.

Mary J Goldman, Brian Craft, Mim Hastie, Kristupas Repečka, Fran McDade, Akhil Kamath, Ayan Banerjee, Yunhai Luo, Dave Rogers, Angela N Brooks, et al. Visualizing and interpreting cancer genomics data via the xena platform. *Nature biotechnology*, 38(6):675–678, 2020.

International Cancer Genome Consortium et al. International network of cancer genome projects. *Nature*, 464(7291):993, 2010.

GenomeOC. Therapeutically applicable research to generate effective treatments, Mar 2021. URL `https://ocg.cancer.gov/programs/target`.

Ethan Cerami, Jianjiong Gao, Ugur Dogrusoz, Benjamin E Gross, Selcuk Onur Sumer, Bülent Arman Aksoy, Anders Jacobsen, Caitlin J Byrne, Michael L Heuer, Erik Larsson, et al. The cbio cancer genomics portal: an open platform for exploring multidimensional cancer genomics data, 2012.

Jianjiong Gao, Bülent Arman Aksoy, Ugur Dogrusoz, Gideon Dresdner, Benjamin Gross, S Onur Sumer, Yichao Sun, Anders Jacobsen, Rileen Sinha, Erik Larsson, et al. Integrative analysis of complex cancer genomics and clinical profiles using the cbioportal. *Science signaling*, 6(269):pl1–pl1, 2013.

Christina Curtis, Sohrab P Shah, Suet-Feung Chin, Gulisa Turashvili, Oscar M Rueda, Mark J Dunning, Doug Speed, Andy G Lynch, Shamith Samarajiwa, Yinyin Yuan, et al. The genomic and transcriptomic architecture of 2,000 breast tumours reveals novel subgroups. *Nature*, 486(7403):346–352, 2012.

Laura Antolini, Patrizia Boracchi, and Elia Biganzoli. A time-dependent discrimination index for survival data. *Statistics in medicine*, 24(24):3927–3944, 2005.

Erika Graf, Claudia Schmoor, Willi Sauerbrei, and Martin Schumacher. Assessment and comparison of prognostic classification schemes for survival data. *Statistics in medicine*, 18(17-18):2529–2545, 1999.

Humza Haider, Bret Hoehn, Sarah Davis, and Russell Greiner. Effective ways to build and evaluate individual survival distributions. *J. Mach. Learn. Res.*, 21(85):1–63, 2020.

Moritz Herrmann, Philipp Probst, Roman Hornung, Vindi Jurinovic, and Anne-Laure Boulesteix. Large-scale benchmark study of survival prediction methods using multi-omics data. *Briefings in bioinformatics*, 22(3):bbaa167, 2021.

Frank E Harrell, Robert M Califf, David B Pryor, Kerry L Lee, and Robert A Rosati. Evaluating the yield of medical tests. *Jama*, 247(18):2543–2546, 1982.

Hajime Uno, Tianxi Cai, Michael J Pencina, Ralph B D'Agostino, and Lee-Jen Wei. On the c-statistics for evaluating overall adequacy of risk prediction procedures with censored survival data. *Statistics in medicine*, 30(10):1105–1117, 2011.

Raphael Sonabend, Andreas Bender, and Sebastian Vollmer. Avoiding c-hacking when evaluating survival distribution predictions with discrimination measures. *Bioinformatics*, 38(17):4178–4184, 2022.

Håvard Kvamme, Ørnulf Borgan, and Ida Scheel. Time-to-event prediction with neural networks and cox regression. *arXiv preprint arXiv:1907.00825*, 2019.

Glenn W Brier et al. Verification of forecasts expressed in terms of probability. *Monthly weather review*, 78(1):1–3, 1950.

Simon Klau, Vindi Jurinovic, Roman Hornung, Tobias Herold, and Anne-Laure Boulesteix. Priority-lasso: a simple hierarchical approach to the prediction of clinical outcome using multi-omics data. *BMC bioinformatics*, 19(1):1–14, 2018.

Qixian Zhong, Jonas W Mueller, and Jane-Ling Wang. Deep extended hazard models for survival analysis. In M. Ranzato, A. Beygelzimer, Y. Dauphin, P.S. Liang, and J. Wortman Vaughan, editors, *Advances in Neural Information Processing Systems*, volume 34, pages 15111–15124. Curran Associates, Inc., 2021. URL https://proceedings.neurips.cc/paper/2021/file/7f6caf1f0ba788cd7953d817724c2b6e-Paper.pdf.

Michel Lang, Martin Binder, Jakob Richter, Patrick Schratz, Florian Pfisterer, Stefan Coors, Quay Au, Giuseppe Casalicchio, Lars Kotthoff, and Bernd Bischl. mlr3: A modern object-oriented machine learning framework in r. *Journal of Open Source Software*, 4(44):1903, 2019.

Raphael Sonabend, Franz J Király, Andreas Bender, Bernd Bischl, and Michel Lang. mlr3proba: An r package for machine learning in survival analysis. *Bioinformatics*, 37(17):2789–2791, 2021.

Kevin Ushey and Hadley Wickham. *renv: Project Environments*, 2024. URL https://rstudio.github.io/renv/. R package version 1.0.5, https://github.com/rstudio/renv.

Jerome Friedman, Trevor Hastie, and Robert Tibshirani. Regularization paths for generalized linear models via coordinate descent. *Journal of Statistical Software*, 33(1):1–22, 2010. doi: 10.18637/jss.v033.i01. URL https://www.jstatsoft.org/v33/i01/.

Noah Simon, Jerome Friedman, Trevor Hastie, and Rob Tibshirani. Regularization paths for cox's proportional hazards model via coordinate descent. *Journal of Statistical Software*, 39(5):1–13, 2011. doi: 10.18637/jss.v039.i05. URL https://www.jstatsoft.org/v39/i05/.

Roman Hornung, Frederik Ludwigs, Jonas Hagenberg, and Anne-Laure Boulesteix. Prediction approaches for partly missing multi-omics covariate data: A literature review and an empirical comparison study. *Wiley Interdisciplinary Reviews: Computational Statistics*, 16(1):e1626, 2024.

Adam Paszke, Sam Gross, Francisco Massa, Adam Lerer, James Bradbury, Gregory Chanan, Trevor Killeen, Zeming Lin, Natalia Gimelshein, Luca Antiga, Alban Desmaison, Andreas Kopf, Edward Yang, Zachary DeVito, Martin Raison, Alykhan Tejani, Sasank Chilamkurthy, Benoit Steiner, Lu Fang, Junjie Bai, and Soumith Chintala. Pytorch: An imperative style, high-performance deep learning library. In H. Wallach, H. Larochelle, A. Beygelzimer, F. d'Alché-Buc, E. Fox, and R. Garnett, editors, *Advances in Neural Information Processing Systems 32*, pages 8024–8035. Curran Associates, Inc., 2019. URL http://papers.neurips.cc/paper/9015-pytorch-an-imperative-style-high-performance-deep-learning-library.pdf.

Marian Tietz, Thomas J. Fan, Daniel Nouri, Benjamin Bossan, and skorch Developers. *skorch: A scikit-learn compatible neural network library that wraps PyTorch*, July 2017. URL https://skorch.readthedocs.io/en/stable/.

Sergey Ioffe and Christian Szegedy. Batch normalization: Accelerating deep network training by reducing internal covariate shift. In *International conference on machine learning*, pages 448–456. PMLR, 2015.

Vinod Nair and Geoffrey E Hinton. Rectified linear units improve restricted boltzmann machines. In *ICML*, 2010.

Ilya Loshchilov and Frank Hutter. Decoupled weight decay regularization. In *International Conference on Learning Representations*, 2018.

David Wissel, Daniel Rowson, and Valentina Boeva. Systematic comparison of multi-omics survival models reveals a widespread lack of noise resistance. *Cell Reports Methods*, 3(4), 2023.

Fabian Pedregosa, Gaël Varoquaux, Alexandre Gramfort, Vincent Michel, Bertrand Thirion, Olivier Grisel, Mathieu Blondel, Peter Prettenhofer, Ron Weiss, Vincent Dubourg, et al. Scikit-learn: Machine learning in python. *the Journal of machine Learning research*, 12:2825–2830, 2011.

Sebastian Pölsterl. scikit-survival: A library for time-to-event analysis built on top of scikit-learn. *J. Mach. Learn. Res.*, 21(212):1–6, 2020.
